# The E3 ligase NEDD4L prevents colorectal cancer liver metastasis via degradation of PRMT5 to inhibit the AKT/mTOR signaling pathway

**DOI:** 10.1101/2024.10.21.619337

**Authors:** Zhewen Dong, Xiaofei She, Junxian Ma, Qian Chen, Yaqun Gao, Ruiyan Chen, Huanlong Qin, Hua Gao

**Author notes:** Zhewen Dong, Xiaofei She, Junxian Ma, Qian Chen, and Yaqun Gao contributed equally to this work. Correspondence: Hua Gao.

## Abstract

Colorectal cancer is the second most common cause of cancer mortality worldwide, and liver metastasis is the major cause of death in patients with colorectal cancer. Dysfunctional E3 ligase activity has recently been shown to be associated with colorectal cancer. However, the key E3 ligase affecting colorectal cancer liver metastasis remains unknown. Therefore, we used an shRNA library targeting 156 E3 ubiquitin ligases to perform an *in vivo* loss-of-function screen of a human colorectal cancer cell line in a mouse model of colorectal cancer liver metastasis. The screen revealed that NEDD4L knockdown promoted colorectal cancer liver metastasis. Moreover, overexpression of NEDD4L prevented colorectal cancer liver metastasis. Mechanistic studies revealed that NEDD4L bound to the PPNAY motif in PRMT5 and ubiquitinated PRMT5 to promote its degradation. PRMT5 degradation attenuated the methylation of an arginine residue in AKT1 to inhibit the AKT/mTOR signaling pathway, consequently decreasing colorectal cancer cell proliferation. We are the first to show that PRMT5 is a substrate protein of the E3 ligase NEDD4L and reveals not only the metastasis-inhibiting function of NEDD4L but also a novel mechanism by which NEDD4L suppresses colorectal cancer liver metastasis. These findings may provide a new preventive strategy for liver metastasis.

## 1. Introduction

Colorectal cancer is the second most common cause of cancer mortality worldwide, responsible for approximately 1.5-2 million deaths annually ^[1]^. More than 50% of patients with colorectal cancer develop liver metastasis, which is the major cause of death in patients with colorectal cancer ^[2]^. Ubiquitination is a posttranslational modification ^[3]^. E3 ubiquitin ligases, the enzymes that mediate the essential step of ubiquitination, specifically link the ubiquitin protein to substrate proteins. To date, more than 600 E3 ligases have been identified ^[4,5]^. Recently, various reports have shown that dysfunctional activity of E3 ligases is associated with cancer ^[6–10]^. However, the key E3 ligase affecting colorectal cancer liver metastasis remains unknown.

Neural precursor cell expressed developmentally down-regulated gene 4-like (NEDD4L), a member of the NEDD4 family, is a highly conserved HECT (homologous to E6AP C terminus) E3 ubiquitin ligase ^[11]^. NEDD4L can regulate the abundance of proteins that participate in epithelial Na^+^ channel regulation, DNA repair, autophagy, and antiviral immunity in a manner depending mainly on the substrate proteins ^[12–19]^. Although several studies have revealed the downregulation of NEDD4L in colorectal cancer and suggested that NEDD4L is a tumor suppressor ^[12,14,18,20]^, the role and mechanism of NEDD4L in colorectal cancer liver metastasis have not been elucidated.

Protein arginine methylation, which is catalyzed by protein arginine methyltransferases (PRMTs), is involved in diverse cellular functions, including cell proliferation and differentiation, RNA splicing, and signal transduction ^[21,22]^. Protein arginine methyltransferase 5 (PRMT5), a type II PRMT, catalyzes the formation of monomethylarginine (MMA) and symmetric dimethylarginine (SDMA) in target proteins ^[23]^. Accumulating evidence indicates that PRMT5 is upregulated in colorectal cancer and is thus considered an oncogene ^[24–26]^. Therefore, the protein stability of PRMT5 is important for its function. However, whether the ubiquitin-proteasome system regulates PRMT5 stability is unknown, and the E3 ligase mediates PRMT5 degradation has not been identified. Furthermore, whether the ubiquitination and degradation of PRMT5 affect colorectal cancer liver metastasis is unclear.

Here, we selected 794 short hairpin RNAs (shRNAs) targeting 156 E3 ubiquitin ligases to construct an shRNA library (Table S1, Supporting Information) and used the shRNA library to perform an *in vivo* loss-of-function screen of the human colorectal cancer cell line HCT-15. We focused on metastatic colonization in the liver, the rate-limiting step of the metastatic cascade ^[27]^. Hence, we performed intrasplenic injection of colorectal cancer cells to establish the mouse model of colorectal cancer liver metastasis for the *in vivo* functional screen. The screening results revealed that NEDD4L knockdown promoted colorectal cancer liver metastasis. Moreover, we found that overexpression of NEDD4L prevented colorectal cancer liver metastasis, while an E3 ligase activity-dead mutant of NEDD4L failed to suppress liver metastasis, indicating that the function of NEDD4L is dependent on its E3 ubiquitin ligase activity. To identify the substrate proteins of NEDD4L, we pulled down NEDD4L and used mass spectrometry to analyze the immunoprecipitates. We discovered that PRMT5 is a substrate protein of NEDD4L. Mechanistically, NEDD4L-mediated ubiquitination of PRMT5 promotes its degradation to attenuate the methylarginine of AKT1. This decrease in the methylarginine level of AKT1 leads to inhibition of the AKT/mTOR signaling pathway and a consequent decrease in colorectal cancer cell proliferation. Taken together, the results of our study reveal that NEDD4L decreases the proliferation of colorectal cancer cells, in turn preventing colorectal cancer liver metastasis. This effect relies on NEDD4L-induced ubiquitination and degradation of PRMT5 to inhibit the AKT/mTOR signaling pathway.

## 2. Results

### 2.1. Identification of NEDD4L as a repressor of colorectal cancer liver metastasis through an *in vivo* functional screen

To identify the key E3 ubiquitin ligase that prevents colorectal cancer liver metastasis, we selected 794 shRNAs targeting 156 cancer-related E3 ligases to construct an shRNA library (Table S1, Supporting Information) and used the shRNA library to perform an *in vivo* loss-of-function screen of the human colorectal cancer cell line HCT-15 (Figure 1A). We focused on metastatic colonization in the liver, the rate-limiting step of the metastatic cascade ^[27]^. Hence, we performed intrasplenic injection of colorectal cancer cells to establish the mouse model of colorectal cancer liver metastasis for the *in vivo* functional screen (Figure 1A).

**Figure 1.**
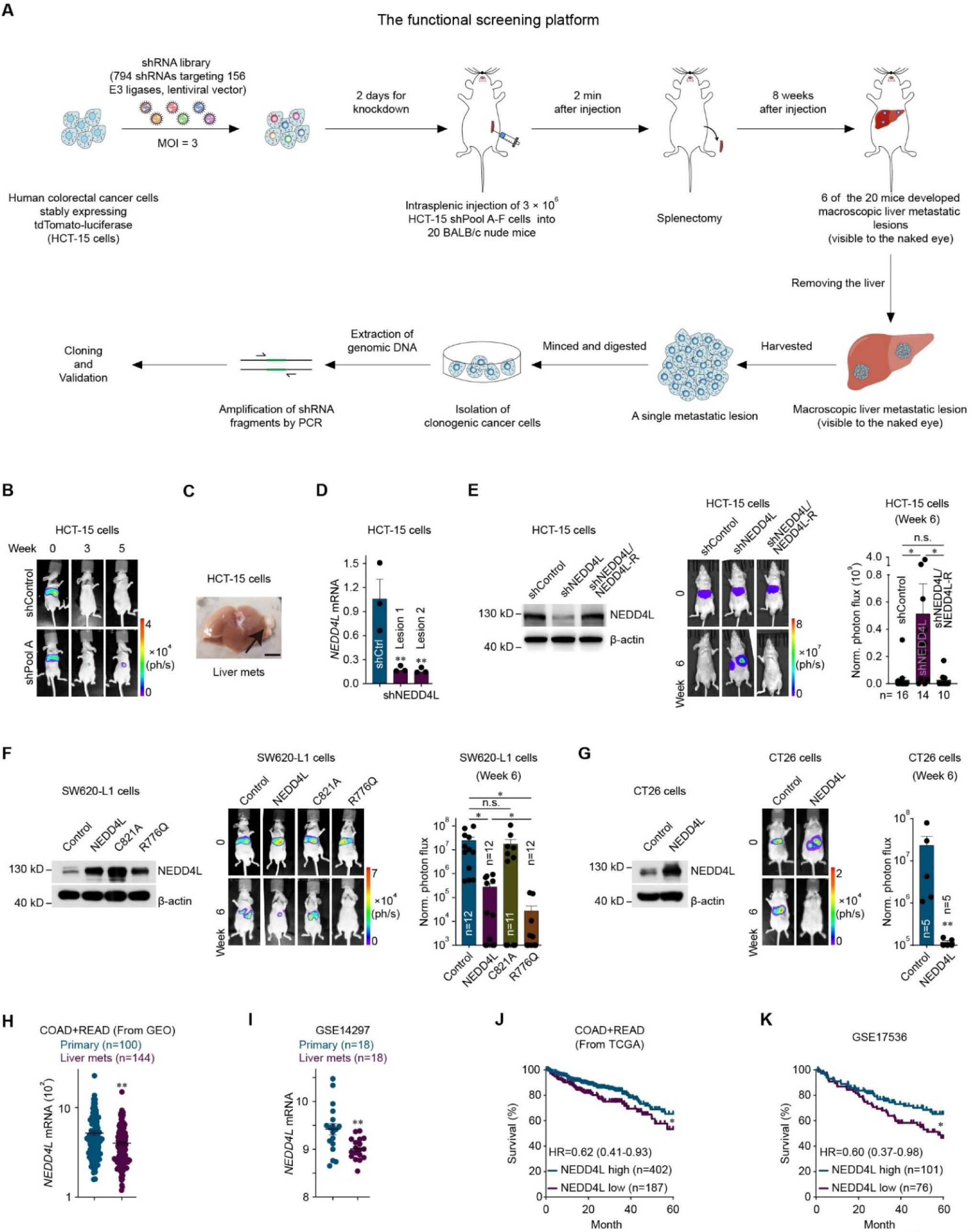
Identification of NEDD4L as a repressor of colorectal cancer liver metastasis through an *in vivo* functional screen. **(A)** Schematic of the loss-of-function screening strategy based on an shRNA library (containing 794 shRNAs targeting 156 E3 ubiquitin ligases) used to identify repressor genes of colorectal cancer liver metastasis. HCT-15 human colorectal cancer cells stably expressing tdTomato-luciferase were transduced independently with each of the six subpools at a multiplicity of infection (MOI) of 3:1. The transduced HCT-15 cells (3 × 10^6^ cells/100 μl of PBS) were subsequently implanted into BALB/c nude mice via intrasplenic injection 2 days later. Eight weeks after injection, macroscopic metastatic lesions in the liver (visible to the naked eye) were harvested and minced to isolate cancer cells. The clonogenic cancer cells were expanded in medium supplemented with puromycin, and the genomic DNA of the cancer cells was extracted. The resident shRNA-encoding sequences in the genomic DNA were amplified with primers, the PCR products were subsequently cloned and inserted into a TA cloning vector, and were analyzed via Sanger sequencing. The mRNA levels of the shRNAs targeting E3 ubiquitin ligases were analyzed by qPCR to assess the on-target knockdown efficiency of each shRNA. MOI, multiplicity of infection. (**B)** Representative bioluminescence images of BALB/c nude mice implanted with parental HCT-15 human colorectal cancer cells or HCT-15 human colorectal cancer cells transduced with the shRNA library targeting E3 ubiquitin ligases (3 × 10^6^ cells; shPool A containing shNEDD4L). (**C)** Representative screening results for liver metastatic lesions in BALB/c nude mice implanted with HCT-15 human colorectal cancer cells transduced with the shRNA library targeting human E3 ubiquitin ligases (3 × 10^6^ cells; shPool A containing shNEDD4L). The arrow indicates liver lesions (macroscopically visible). Scale bar, 1 cm. (**D)** mRNA level of NEDD4L in HCT-15 cells recovered from two liver metastatic lesions containing shNEDD4L that were formed by HCT-15 cells transduced with shPool A. (**E)** Representative western blots showing NEDD4L expression in control HCT-15 cells (shControl), NEDD4L-knockdown HCT-15 cells (shNEDD4L) and HCT-15 cells with knockdown of NEDD4L and restoration with wild-type NEDD4L resistant to shNEDD4L targeting (shNEDD4L/NEDD4L-R) (left). Bioluminescence imaging results (middle) and quantification of liver metastases (right) in BALB/c nude mice implanted with shControl, shNEDD4L or shNEDD4L/NEDD4L-R HCT-15 cells (3 × 10^6^ cells) via intrasplenic injection. The n-values denote the number of mice per group, and three independent western blot analyses were performed. (**F)** Representative western blots showing NEDD4L and NEDD4L mutant expression in control SW620-L1 cells (Control), and SW620-L1 cells overexpressing wild-type NEDD4L (NEDD4L), an E3 ligase activity-dead mutant of NEDD4L (C821A) or a constitutively active mutant of NEDD4L (R776Q) (left). Bioluminescence imaging results (middle) and quantification of liver metastases (right) of BALB/c nude mice implanted with Control, NEDD4L, C821A or R776Q SW620-L1 cells (1 × 10^6^ cells) via intrasplenic injection. To induce the overexpression of NEDD4L or its mutants in these cells, doxycycline (100 μg/mouse) was administered intraperitoneally on the day of cancer cell injection (day 0) and was then administered orally (400 ppm in chow combined with 2 mg/ml in water) from day 0 to the experimental endpoint. The n-values denote the number of mice per group, and three independent western blot analyses were performed. (**G)** Representative western blots showing NEDD4L expression in control CT26 cells (Control) and CT26 cells overexpressing wild-type NEDD4L (NEDD4L) (left). Bioluminescence imaging results (middle) and quantification of liver metastases (right) of BALB/c mice implanted with Control or NEDD4L CT26 cells (3 × 10^5^ cells) via intrasplenic injection. To induce the overexpression of NEDD4L in the cells, doxycycline (100 μg/mouse) was administered intraperitoneally on the day of cancer cells injection (day 0) and was then administered orally (400 ppm in chow in combination with 2 mg/ml in water) from day 0 to the experimental endpoint. The n-values denote the number of mice per group, and three independent western blot analyses were performed. (**H)** NEDD4L mRNA expression in the primary tumors and unpaired liver metastatic lesions of patients with colorectal cancer in the combined dataset from GLP570 platform (Affymetrix Human Genome U133 Plus 2.0 Array). The combined dataset consists of GSE10961, GSE18462, GSE28702, GSE40367 and GSE41568. The n values denote the number of patients per group. (**I)** NEDD4L mRNA expression in the primary tumors and paired liver metastatic lesions of patients with colorectal cancer in GSE14297 dataset. The n values denote the number of patients per group. (**J)** Kaplan-Meier curves of 5-year survival for patients with colorectal cancer represented in TCGA datasets (COAD and READ, n = 589) stratified by the NEDD4L mRNA expression level. HR, hazard ratio. (**K)** Kaplan-Meier curves of 5-year survival for patients with colorectal cancer represented in GSE17536 dataset (n = 177) stratified by the NEDD4L mRNA expression level. HR, hazard ratio. The data are presented as the mean ± s.e.m. values. *P*-values were determined by unpaired one-way ANOVA with uncorrected Fisher’s LSD test (D, E, and F), unpaired two-tailed Student’s t-test with Welch’s correction (G), the Wilcoxon rank-sum test (H and I), or the log-rank test (J and K). * *P* < 0.05; ** *P* < 0.01; n.s., not significant.

In the screen, shNEDD4L (TRCN0000000905), targeting NEDD4L was identified in 2 independent liver metastatic lesions in two mice and was the top-ranked shRNA (Figure 1B and C and Figure S1A, Supporting Information). NEDD4L knockdown in HCT-15 cells recovered from the 2 liver metastatic lesions was confirmed at the transcriptional level (Figure 1D). Notably, the shNEDD4L had the highest knockdown efficiency among the 5 shRNAs in the library, and only shNEDD4L (TRCN0000000905) was identified.

Consistent with the screening results, in the validation experiment, knockdown of NEDD4L with shNEDD4L (TRCN0000000905) promoted the liver metastasis of HCT-15 cells (Figure 1E). To prevent any off-target effect of shNEDD4L, NEDD4L-R, which was resistant to shNEDD4L targeting, was used to restore NEDD4L expression in NEDD4L-knockdown colorectal cancer cells, since we did not find in our system that the knockdown efficiency of any published shRNA (different from the 5 shRNAs in the library) was similar to that of shNEDD4L (TRCN0000000905). In the functional rescue experiment, overexpression of NEDD4L-R in NEDD4L-knockdown colorectal cancer cells prevented the liver metastasis of these cells, suggesting that restoring NEDD4L expression reversed the effect of NEDD4L knockdown on colorectal cancer liver metastasis (Figure 1E). Moreover, in SW620-L1 and CT26 cells, which are highly metastatic colorectal cancer cell lines derived from humans and mice, respectively, overexpression of wild-type NEDD4L prevented the liver metastasis of cancer cells (Figure 1F and G and Figure S1B, Supporting Information).

To determine whether the E3 ligase activity of NEDD4L plays a critical role in its ability to inhibit metastasis, an E3 ligase activity-dead mutant of NEDD4L (NEDD4L C821A) and a constitutively active mutant of NEDD4L (NEDD4L R776Q) ^[28]^ were generated and overexpressed in SW620-L1 cells (Figure 1F and Figure S1C, Supporting Information). NEDD4L C821A overexpression failed to prevent colorectal cancer liver metastasis, and NEDD4L R776Q overexpression prevented colorectal cancer liver metastasis more potently than NEDD4L overexpression did (Figure 1F), suggesting that the E3 ligase activity of NEDD4L is important for its ability to inhibit metastasis.

Additionally, in our analysis of clinical samples, although NEDD4L expression was high in primary tumors, it was lower in liver metastatic (Figure 1H and I). Moreover, low expression of NEDD4L in primary tumors was correlated with poor 5-year overall survival outcomes in patients with colorectal cancer (Figure 1J and K).

### 2.2. NEDD4L decreases cell proliferation to prevent colorectal cancer liver metastasis through inhibition of the AKT/mTOR signaling pathway

The function of NEDD4L related to metastatic colonization was explored since colorectal cancer cells were injected directly into the bloodstream. The *in vitro* results revealed that the proliferation of colorectal cancer cells was suppressed by NEDD4L, although the epithelial-mesenchymal transition (EMT), invasion and stemness of the colorectal cancer cells were not affected (Figure 2A and B and Figure S2A-E, Supporting Information). The proliferation of colorectal cancer cells stimulated with amino acids (AA) and insulin was subsequently examined since the liver is the main organ for storing AA and insulin ^[29,30]^. The proliferation of colorectal cancer cells stimulated with AA and insulin was also suppressed by NEDD4L (Figure 2C and D). However, overexpression of NEDD4L C821A (an E3 ligase activity-dead mutant) failed to decrease the proliferation of colorectal cancer cells, and the overexpression of NEDD4L R776Q (a constitutively active mutant) increased the suppressive effect of NEDD4L on the proliferation of colorectal cancer cells (Figure 2B and D). Moreover, the *in vivo* results revealed that the overexpression of NEDD4L decreased the proliferation of liver-metastatic SW620-L1 human colorectal cancer cells (Figure 2E).

**Figure 2.**
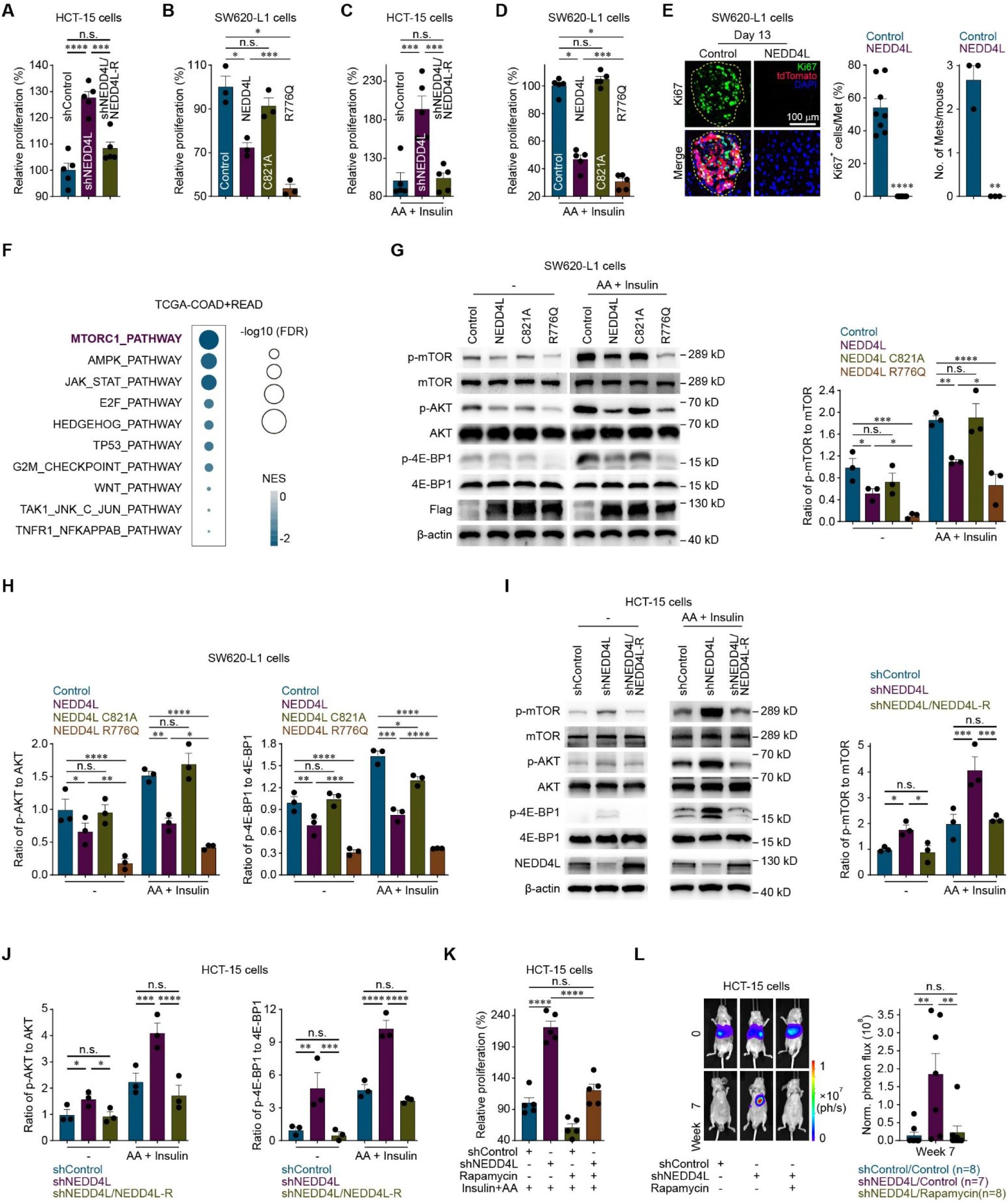
NEDD4L decreases cell proliferation to prevent colorectal cancer liver metastasis through inhibition of the AKT/mTOR signaling pathway. (**A)** *In vitro* proliferation assay of control HCT-15 cells (shControl), NEDD4L-knockdown HCT-15 cells (shNEDD4L) and HCT-15 cells with NEDD4L knockdown and restoration with wild-type NEDD4L resistant to shRNA targeting NEDD4L (shNEDD4L/NEDD4L-R) (1,000 cells) cultured for 72 hr. Five independent CCK-8 assays were performed. (**B)** *In vitro* proliferation assay of control SW620-L1 cells (Control) and SW620-L1 cells with overexpression of NEDD4L wild-type (NEDD4L), an E3 ligase activity-dead mutant of NEDD4L (C821A) or a constitutively active mutant of NEDD4L (R776Q) (3,000 cells) induced by treatment with 2 µg/ml doxycycline and cultured for 72 hr. Three independent CCK-8 assays were performed. (**C)** *In vitro* proliferation assay of control HCT-15 cells (shControl), NEDD4L-knockdown HCT-15 cells (shNEDD4L) and HCT-15 cells with NEDD4L knockdown and restoration with wild-type NEDD4L resistant to shRNA targeting NEDD4L (shNEDD4L/NEDD4L-R) (1,000 cells) cultured with 200 μM amino acids (AA) and 800 nM insulin for 24 hr. Five independent CCK-8 assays were performed. (**D)** *In vitro* proliferation assay of control SW620-L1 cells (Control) and SW620-L1 cells with overexpression of NEDD4L wildt-type (NEDD4L), an E3 ligase activity-dead mutant of NEDD4L (C821A) or a constitutively active mutant of NEDD4L (R776Q) (3,000 cells) induced by treatment with 2 µg/ml doxycycline and cultured with 200 μM AA and 800 nM insulin for 24 hr. Five independent CCK-8 assays were performed. (**E)** Representive images (right), quantification of Ki67 (green)-positive SW620-L1 cells (tdTomato) in the liver metastatic lesions of SW620-L1 cell-implanted BALB/c nude mice (middle), and quantification of liver metastatic lesions (right) on day 13 after intrasplenic injection of control or NEDD4L-overexpressing SW620-L1 cells (1 × 10^6^ cells). To induce the overexpression of NEDD4L in the cells, doxycycline (100 μg/mouse) was administered intraperitoneally on the day 7 after cancer cells injection and was then administered orally (400 ppm in chow in combination with 2 mg/ml in water) from day 7 to day 13 after cancer cell injection. The areas with staining of Ki67^+^ cancer cells and tdTomato^+^ cancer cells in the ROI were calculated via ImageJ. Three random sections were used to calculate the average percentage of Ki67^+^ cancer cells per liver metastatic lesion (8 liver metastatic lesions in 3 mice in the control group and 0 liver metastatic lesions in 3 mice in the NEDD4L group). Yellow dotted lines (ROIs) or Met, liver metastatic lesions; ROI, region of interest (**F)** GSEA revealed an association between NEDD4L expression and the enrichment of gene in proliferation-related signaling pathways in clinical colorectal cancer samples from the TCGA datasets (COAD and READ, n = 589). GSEA, gene set enrichment analysis. NES, normalized enrichment score. FDR, false discovery rate. (**G-H)** Representative western blots (G, left) and quantification of p-mTOR levels (normalized to mTOR levels, G, right), p-AKT levels (normalized to AKT levels, H, right), and p-4E-BP1 levels (normalized to 4E-BP1 levels, H, right) in control SW620-L1 cells (Control) and SW620-L1 cells with overexpression of wild-type NEDD4L (NEDD4L), an E3 ligase activity-dead mutant of NEDD4L (C821A) or a constitutively active mutant of NEDD4L (R776Q) induced by treatment with 2 µg/ml doxycycline (for 24 hr) and incubated with or without 200 μM AA (for 15 min) and 800 nM insulin (for 10 min). Three independent western blot analyses were performed. (**I, J)** Representative western blots (I, left) and quantification of p-mTOR levels (normalized to mTOR levels, I, right), p-AKT levels (normalized to AKT levels, J, left), and p-4E-BP1 levels (normalized to 4E-BP1 levels, J, right) in control HCT-15 cells (shControl), NEDD4L-knockdown HCT-15 cells (shNEDD4L) and HCT-15 cells with knockdown of NEDD4L and restoration with wild-type NEDD4L resistant to shRNA targeting NEDD4L (shNEDD4L/NEDD4L-R) incubated with or without 200 μM AA (for 15 min) and 800 nM insulin (for 10 min). Three independent western blot analyses were performed. (**K)** *In vitro* proliferation assay of control HCT-15 cells (shControl) and NEDD4L-knockdown HCT-15 cells (shNEDD4L) (1,000 cells) cultured with or without 100 nM rapamycin in combination with 200 μM AA and 800 nM insulin for 24 hr. Five independent CCK-8 assays were performed. (**L)** Bioluminescence imaging results (left) and quantification of liver metastases (right) in BALB/c nude mice implanted with control HCT-15 cells (shControl) or NEDD4L-knockdown HCT-15 cells (shNEDD4L) (3 × 10^6^ cells) via intrasplenic injection. Rapamycin (2 mg/kg) was administered intraperitoneally every two days from the day of cancer cell injection (day 0) to the experimental endpoint. The n-values denote the number of mice per group. The data are presented as the mean ± s.e.m. values. *P*-values were determined by unpaired one-way ANOVA with uncorrected Fisher’s LSD test (A, B, C, D, K, and L), unpaired two-tailed Student’s t-test with Welch’s correction (E), or unpaired two-way ANOVA with uncorrected Fisher’s LSD test (G, H, I, and J). * *P* < 0.05; ** *P* < 0.01; *** *P* < 0.001; **** *P* < 0.0001; n.s., not significant.

Gene set enrichment analysis (GSEA) based on colorectal cancer patient data revealed that the activity of the mTOR signaling pathway was negatively correlated with the expression of NEDD4L (Figure 2F). Then, since AA and insulin are the main activators of the AKT/mTOR signaling pathway ^[31,32]^, the effects of NEDD4L on the AKT/mTOR signaling pathway in the context of AA and insulin stimulation were examined. NEDD4L inhibited the AKT/mTOR signaling pathway regardless of stimulation with AA and/or insulin (Figure 2G and H and Figures S2F and G and S3A-F, Supporting Information), whereas NEDD4L did not affect the AKT/mTOR signaling pathway under stimulation with glucose (Figure S3G and H, Supporting Information). In contrast, knockdown of NEDD4L upregulated the AKT/mTOR signaling pathway under both the basal and activating conditions (Figure 2I and J and Figure S4A-F, Supporting Information). The functional rescue experiment revealed that overexpression of NEDD4L-R in NEDD4L-knockdown HCT-15 cells inhibited the AKT/mTOR signaling pathway, suggesting that restoring NEDD4L expression reversed the effect of NEDD4L knockdown on the AKT/mTOR signaling pathway (Figure 2I and J). In addition, overexpression of the E3 ligase activity-dead NEDD4L C821A mutant failed to inhibit the AKT/mTOR signaling pathway, and overexpression of the constitutively active NEDD4L R776Q mutant inhibited the AKT/mTOR signaling pathway more potently than did overexpression of wild-type NEDD4L (Figure 2G and H). These results suggested that NEDD4L inhibited the AKT/mTOR signaling pathway and that this effect was dependent on its E3 ligase activity.

Furthermore, the *in vitro* experiments revealed that the effect of NEDD4L knockdown on cancer cell proliferation was reversed by treatment with rapamycin, which is a selective mTOR inhibitor that suppresses the AKT/mTOR signaling pathway (Figure 2K and Figure S4G, Supporting Information). Consistent with the results of the *in vitro* experiments, the effect of NEDD4L knockdown on the liver metastasis of HCT-15 cells was reversed by rapamycin treatment (Figure 2L). These results suggested that NEDD4L decreased cancer cell proliferation through the inhibition of the AKT/mTOR signaling pathway to prevent the liver metastasis of colorectal cancer cells.

### 2.3. NEDD4L ubiquitinates PRMT5 to promote PRMT5 degradation

As demonstrated above, NEDD4L inhibited the AKT/mTOR signaling pathway in a manner dependent on its E3 ligase activity. Next, we examined whether proteins in the AKT/mTOR signaling pathway are direct substrates of NEDD4L. The results showed that NEDD4L had no effect on the protein level of AKT, TSC2, or Rheb (Figure S5A and B, Supporting Information). To identify the direct substrate proteins, Flag-NEDD4L and HA-ubiquitin were pulled down (Figure 3A and B), and the immunoprecipitates were analyzed by mass spectrometry. The PRMT5-WDR77 complex was identified as 2 top candidate proteins (Figure 3C and D and Table S2, Supporting Information). PRMT5 has been reported to be able to active AKT ^[25,33]^, and the presence of WDR77 as a cofactor can increase the affinity of PRMT5 for its target proteins ^[34]^. The validation experiment revealed that NEDD4L ubiquitinated PRMT5 to promote its degradation (Figure 3E-G and Figure S5C and D, Supporting Information), which was dependent on the E3 ligase activity of NEDD4L (Figure 3E and H). However, NEDD4L did not affect the stability of WDR77 (Figure S5C, Supporting Information), indicating that WDR77 is not a direct substrate protein of NEDD4L.

**Figure 3.**
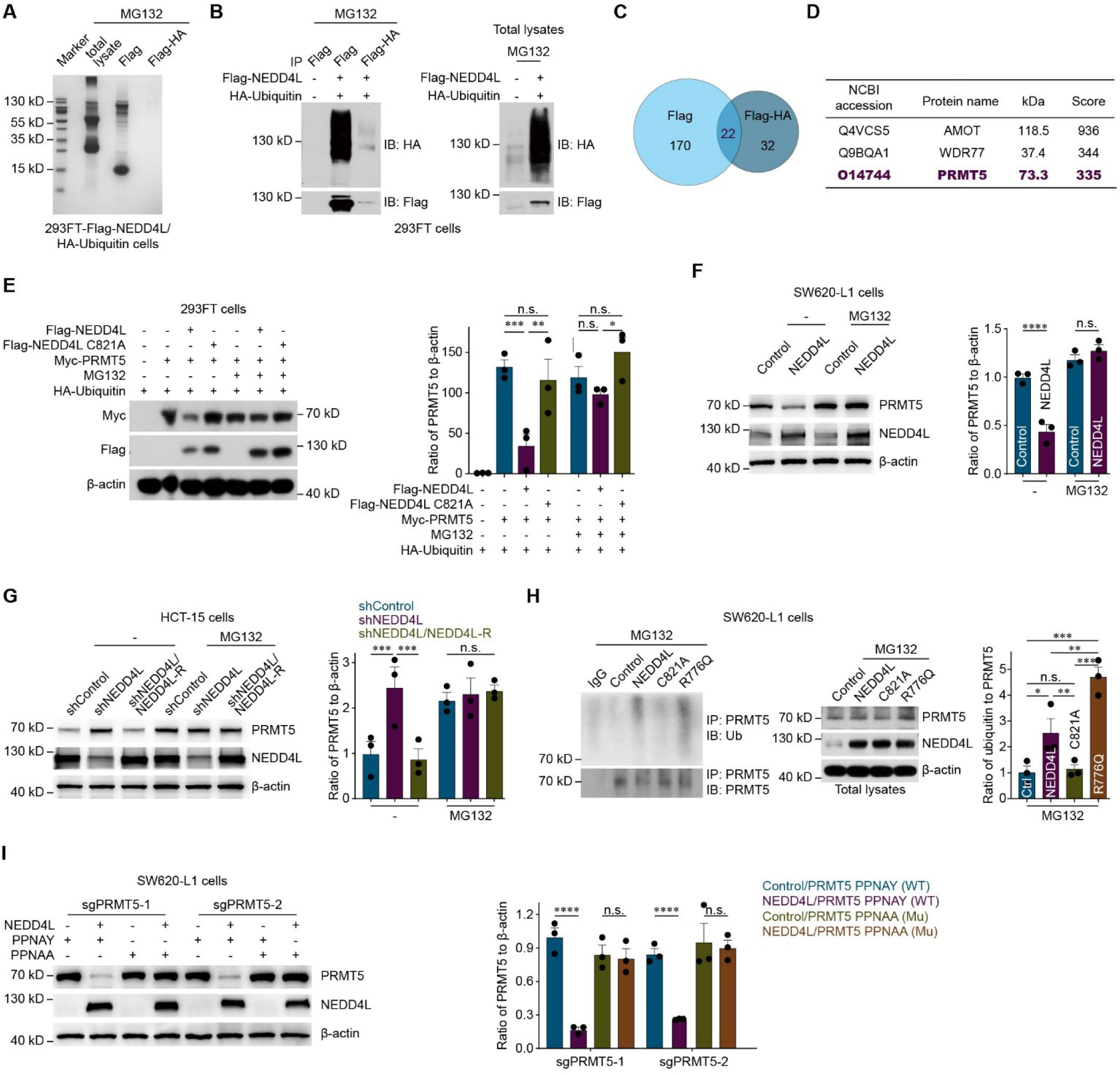
NEDD4L ubiquitinates PRMT5 to promote its degradation. (**A)** EZBlue™ staining of gels containing the total lysate of 293FT cells overexpressing Flag-NEDD4L and HA-ubiquitin. The immunoprecipitates obtained from an immunoprecipitation assay of 293FT cells with overexpression of Flag-NEDD4L and HA-ubiquitin were incubated with 20 μM MG132 for 12 hr and pulled down by Flag or Flag and HA agarose. (**B)** Representative western blots of proteins immunoprecipitated from 293FT cells overexpressing Flag-NEDD4L and HA-ubiquitin overexpression that were incubated with 20 μM MG132 for 12 hr and pulled down by Flag or Flag and HA agarose. Three independent western blot analyses were performed. (**C)** Venn diagram showing the overlap among the candidate proteins identified by mass spectrometry analysis of immunoprecipitates from 293FT cells overexpressing Flag-NEDD4L and HA-ubiquitin that were incubated with 20 μM MG132 for 12 hr and pulled down by Flag or Flag and HA agarose. (**D)** List of the top 3 candidate substrate proteins of NEDD4L identified via the mass spectrometry analysis. (**E)** Representative western blots (left) and quantification of Myc-PRMT5 expression (normalized to β-actin expression, right) in control 293FT cells with HA-ubiquitin overexpression and 293FT cells with HA-ubiquitin overexpression and overexpression of wild-type NEDD4L (Flag-NEDD4L) or an E3 ligase activity-dead mutant of NEDD4L (Flag-NEDD4L C821A) transfected with or without Myc-PRMT5 and incubated with or without 20 μM MG132 for 12 hr. Three independent western blot analyses were performed. (**F)** Representative western blots (left) and quantification of PRMT5 expression (normalized to β-actin expression, right) in control SW620-L1 cells (Control), and SW620-L1 cells overexpressing wild-type NEDD4L (NEDD4L) induced by treatment with 2 µg/ml doxycycline (for 24 hr) and incubated with or without 20 μM MG132 for 12 hr. Three independent western blot analyses were performed. (**G)** Representative western blots (left) and quantification of PRMT5 expression (normalized to β-actin expression, right) in control HCT-15 cells (shControl), NEDD4L-knockdown HCT-15 cells (shNEDD4L) and HCT-15 cells with NEDD4L knockdown and restoration with wild-type NEDD4L resistant to shRNA targeting NEDD4L (shNEDD4L/NEDD4L-R) incubated with or without 20 μM MG132 for 12 hr. Three independent western blot analyses were performed. (**H)** Representative western blots (left) and quantification of PRMT5 ubiquitination (normalized to PRMT5 expression, right) in control SW620-L1 cells (Control) and SW620-L1 cells with overexpression of wild-type NEDD4L (NEDD4L), an E3 ligase activity-dead mutant of NEDD4L (C821A) or a constitutively active mutant of NEDD4L (R776Q) induced by treatment with 2 µg/ml doxycycline (for 24 hr) and incubated with 20 μM MG132 for 12 hr. Three independent western blot analyses were performed. (**I)** Representative western blots (left) and quantification of exogenous Flag-PRMT5 and a NEDD4L binding motif mutant of Flag-PRTM5 (normalized to β-actin expression, right) in control SW620-L1 cells and NEDD4L-overexpressing SW620-L1 cells with knockout of endogenous PRMT5 (sgPRMT5-1 or sgPRMT5-2) and overexpression of PRMT5 containing the wild-type NEDD4L binding motif (PPNAY), or mutant NEDD4L binding motif (PPNAA). To induce the overexpression of NEDD4L in these cells, the cells was induced by treatment with 2 µg/ml doxycycline (for 24 hr). Three independent western blot analyses were performed. The data are presented as the mean ± s.e.m. values. *P*-values were determined by unpaired two-way ANOVA with uncorrected Fisher’s LSD test (E, F, G, and I), or unpaired one-way ANOVA with uncorrected Fisher’s LSD test (H). * *P* < 0.05; ** *P* < 0.01; *** *P* < 0.001; **** *P* < 0.0001; n.s., not significant.

NEDD4L binds to PPxY motifs in its substrate proteins via its WW domain ^[35]^.To confirm the function of the PPNAY motif in PRMT5, motif mutants of PRMT5 (PPNAA and PANAY) were used (Figure S5E, Supporting Information). In cells with knockout of endogenous PRMT5, NEDD4L failed to promote the degradation of either of these PRMT5 mutants (Figure 3I and Figure S5E and F, Supporting Information). These results suggested NEDD4L ubiquitinated PRMT5 to promote its degradation and that this ability was dependent on the PPNAY motif in PRMT5.

### 2.4. PRMT5 activates the AKT/mTOR signaling pathway to increase cell proliferation and promote colorectal cancer liver metastasis

Recent studies have indicated that PRMT5 methylates and activates AKT and consequently increases cancer cell proliferation ^[25,33]^. Consistent with published studies, overexpression of PRMT5 activated the AKT/mTOR signaling pathway (Figure 4A and B and Figure S6A-F, Supporting Information) and consequently increased the proliferation of colorectal cancer cells (Figure 4C), whereas overexpression of PRMT5 R368A (a methyltransferase-inactive mutant of PRMT5) failed to activate the AKT/mTOR signaling pathway and had no effect on colorectal cancer cell proliferation (Figure 4A-C). Both the knockout and knockdown of PRMT5, as well as the inhibition of PRMT5 by EPZ015666, a selective PRMT5 inhibitor that targets the substrate binding pocket of PRMT5 ^[36]^, inhibited the AKT/mTOR signaling pathway (Figure 4D and E and Figure S6G-J, Supporting Information), consequently decreasing the proliferation of colorectal cancer cells (Figure 4F and Figure S6K and L, Supporting Information). Moreover, the effect of PRMT5 on colorectal cancer cell proliferation was reversed by treatment with rapamycin, a selective mTOR inhibitor ^[37]^ that suppresses the AKT/mTOR signaling pathway (Figure 4G-H).

**Figure 4.**
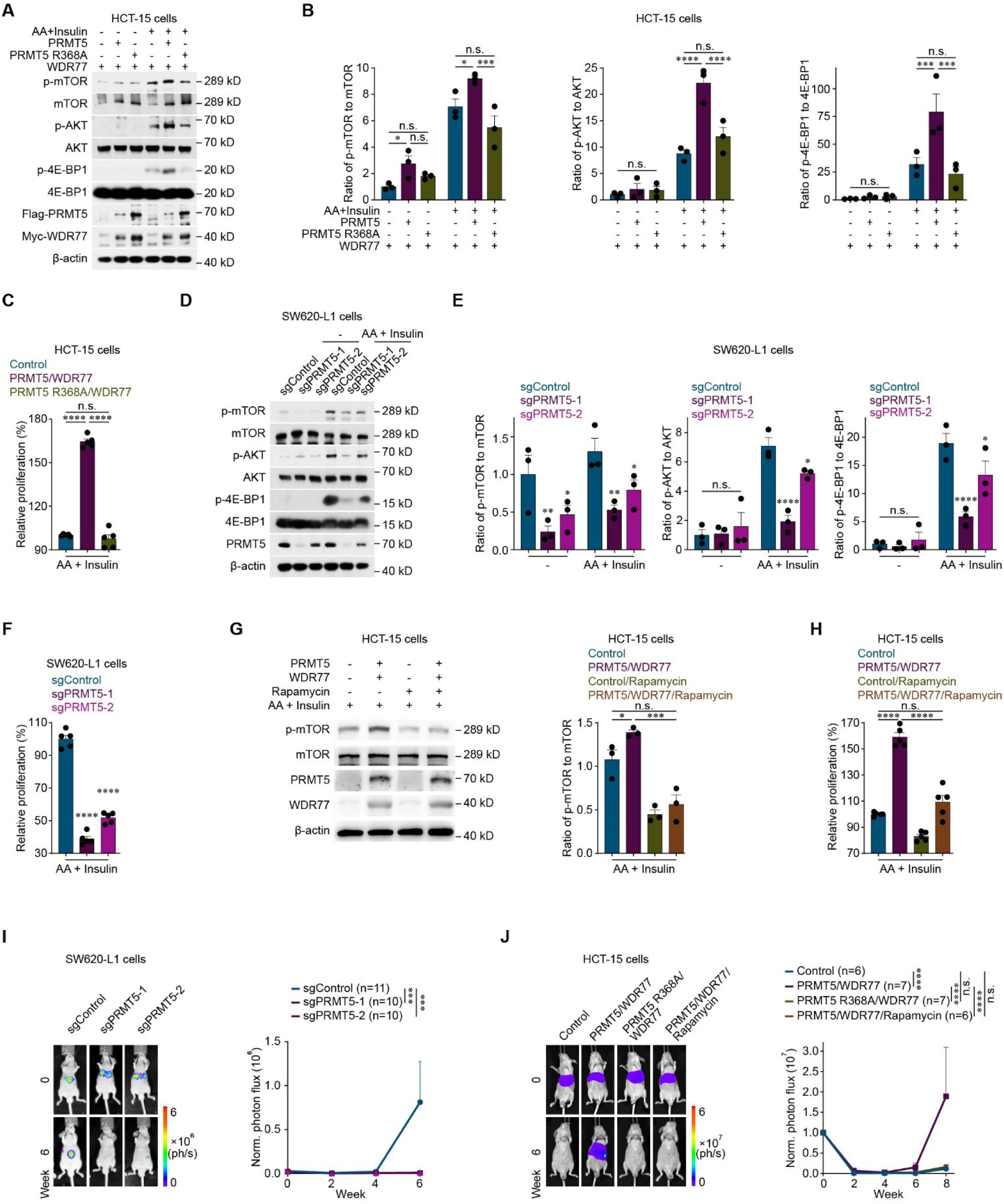
PRMT5 activates the AKT/mTOR signaling pathway to increase cell proliferation and promote colorectal cancer liver metastasis. (**A, B)** Representative western blots (A) and quantification of p-mTOR levels (normalized to mTOR levels, B, left), p-AKT levels (normalized to AKT levels, B, middle), and p-4E-BP1 levels (normalized to 4E-BP1 levels, B, right) in control HCT-15 cells and HCT-15 cells with overexpression of WDR77 in combination with wild-type PRMT5 (PRMT5/WDR77) or a methyltransferase-inactive mutant of PRMT5 (PRMT5 R368A/WDR77) incubated with or without 200 μM amino acids (AA; for 15 min) and 800 nM insulin (for 10 min). Three independent western blot analyses were performed. (**C)** *In vitro* proliferation assay of control HCT-15 cells (Control) and HCT-15 cells with overexpressing WDR77 in combination with wild-type PRMT5 (PRMT5/WDR77) or a methyltransferase-inactive mutant of PRMT5 (PRMT5 R368A/WDR77) (1,000 cells) cultured with 200 μM AA and 800 nM insulin for 24 hr. Five independent CCK-8 assays were performed. (**D, E)** Representative western blots (D, left) and quantification of p-mTOR levels (normalized to mTOR levels, E, left), p-AKT levels (normalized to AKT levels, E, middle), and p-4E-BP1 levels (normalized to 4E-BP1 levels, E, right) in control SW620-L1 cells (sgControl) and PRMT5-knockout SW620-L1 cells (sgPRMT5-1 or sgPRMT5-2) incubated with or without 200 μM AA (for 15 min) and 800 nM insulin (for 10 min). Three independent western blot analyses were performed. (**F)** *In vitro* proliferation assay of control SW620-L1 cells (sgControl) and PRMT5-knockout SW620-L1 cells (sgPRMT5-1 or sgPRMT5-2, 3,000 cells) cultured with 200 μM AA and 800 nM insulin for 24 hr. Five independent CCK-8 assays were performed. (**G)** Representative western blots (left) and quantification of p-mTOR levels (normalized to mTOR levels, right) in control HCT-15 cells and HCT-15 cells overexpressing WDR77 in combination with wild-type PRMT5 (PRMT5/WDR77) cultured with or without 100 nM rapamycin (24 hr) in combination with 200 μM AA (for 15 min) and 800 nM insulin (for 10 min). Three independent western blot analyses were performed. (**H)** *In vitro* proliferation assay of control HCT-15 cells (Control) and HCT-15 cells overexpressing WDR77 in combination with wild-type PRMT5 (PRMT5/WDR77) (1,000 cells) cultured with or without 100 nM rapamycin combined with 200 μM AA and 800 nM insulin for 24 hr. Five independent CCK-8 assays were performed. (**I)** Bioluminescence imaging results (left) and quantification of liver metastases (right) in BALB/c nude mice implanted with control SW620-L1 cells (sgControl) or PRMT5-knockout SW620-L1 cells (sgPRMT5-1 or sgPRMT5-2, 1 × 10^6^ cells) via intrasplenic injection. The n-values denote the number of mice per group. (**J)** Bioluminescence imaging results (left) and quantification of liver metastases (right) in BALB/c nude mice implanted with control HCT-15 cells (Control), HCT-15 cells overexpressing WDR77 in combination with wild-type PRMT5 (PRMT5/WDR77) or HCT-15 cells overexpressing WDR77 in combination with a methyltransferase-inactive mutant of PRMT5 (PRMT5 R368A/WDR77) (3 × 10^6^ cells) via intrasplenic injection. Rapamycin (2 mg/kg) was administered intraperitoneally every two days from the day cancer cell injection (day 0) to the experimental endpoint. The n-values denote the number of mice per group. The data are presented as the mean ± s.e.m. values. *P*-values were determined by unpaired two-way ANOVA with uncorrected Fisher’s LSD test (B, E, I, and J), or unpaired one-way ANOVA with uncorrected Fisher’s LSD test (C, F, G, and H). * *P* < 0.05; *** *P* < 0.001; **** *P* < 0.0001; n.s., not significant.

Furthermore, the *in vivo* functional experiment showed that knockout of PRMT5 prevented colorectal cancer liver metastasis (Figure 4I) and that overexpression of PRTM5 promoted colorectal cancer liver metastasis (Figure 4J), whereas overexpression of the methyltransferase-inactive PRMT5 R368A mutant did not affect liver metastasis (Figure 4J). In addition, the ability of PRMT5 to promote metastasis was reversed by rapamycin treatment (Figure 4J). These results suggested that PRMT5-mediated activation of the AKT/mTOR signaling pathway increased cancer cell proliferation, in turn promoting colorectal cancer liver metastasis, and that this effect was dependent on the methyltransferase activity of PRMT5.

### 2.5. PRMT5 methylates AKT1 at R391 to activate the AKT/mTOR signaling pathway and increase colorectal cancer cell proliferation

Methylation of arginine 391 (R391) in AKT1 has been reported to play a crucial role in AKT activation ^[25]^. Our experiments revealed that PRMT5 interacted with AKT1 (Figure 5A and B) and methylated the arginine residue in AKT1 to activate AKT (Figure 5C and D and Figure S7A, Supporting Information), whereas other proteins in the AKT/mTOR signaling pathway did not interact with PRMT5 (Figure 5A and B).

**Figure 5.**
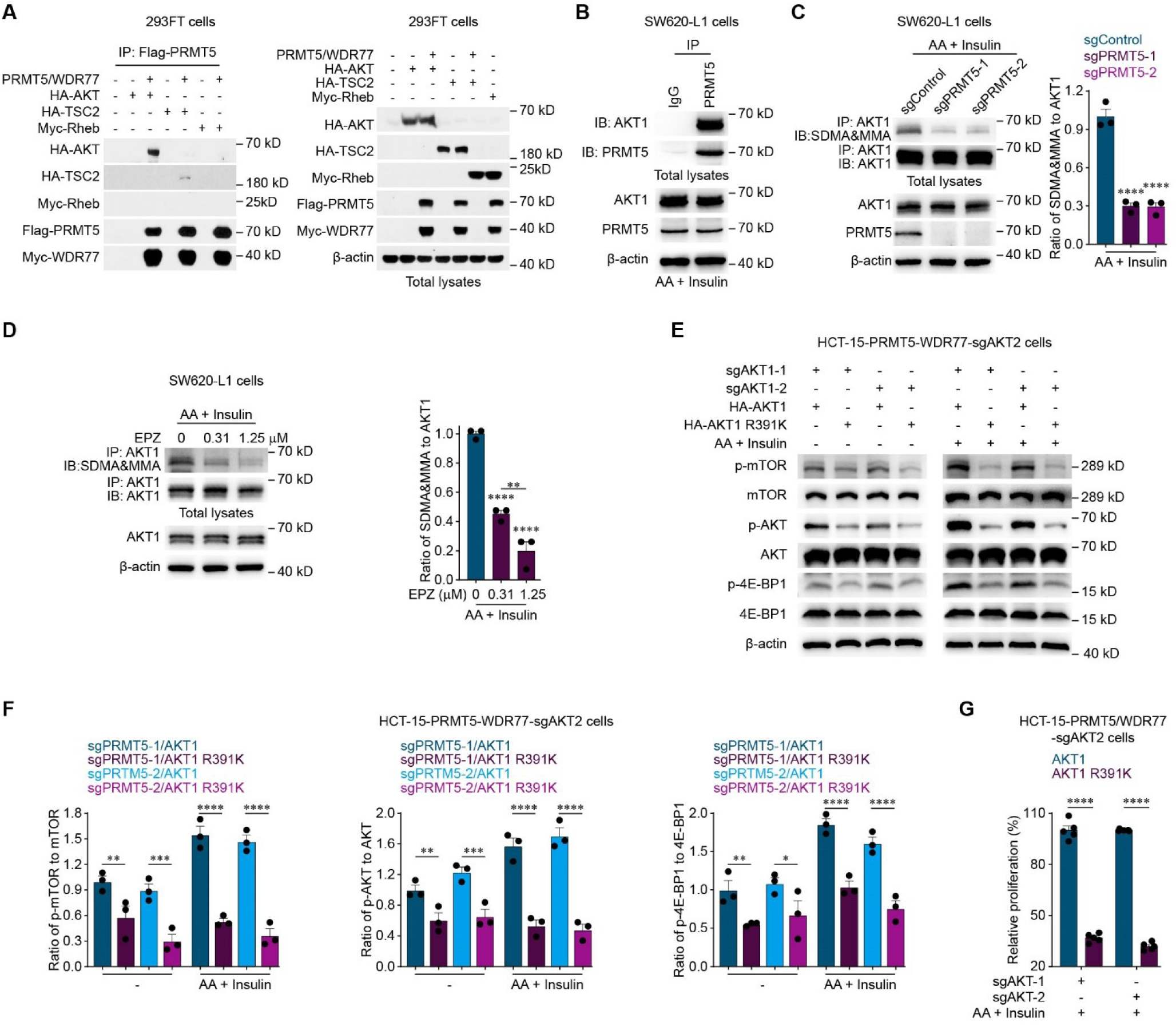
PRMT5 methylates AKT1 R391 to activate the AKT/mTOR signaling pathway and increase colorectal cancer cell proliferation. (**A)** Representative western blots of Flag-PRMT5 immunoprecipitates and total lysates of control 293FT cells and 293FT cells with HA-AKT, HA-TSC2, or Myc-Rheb overexpression with or without Flag-PRMT5 and Myc-WDR77 overexpression. Three independent western blot analyses were performed. (**B)** Representative western blots showing PRMT5 immunoprecipitates and total lysates of SW620-L1 cells incubated with 200 μM amino acids (AA; for 15 min) and 800 nM insulin (for 10 min). Three independent western blot analyses were performed. (**C)** Representative western blots (left) and quantification of AKT1 methylarginine levels (normalized to AKT1 expression, right) in control SW620-L1 cells (sgControl) and PRMT5-knockout SW620-L1 cells (sgPRMT5-1 or sgPRMT5-2) incubated with 200 μM AA (for 15 min) and 800 nM insulin (for 10 min). Three independent western blot analyses were performed. (**D)** Representative western blots (left) and quantification of AKT1 methylarginine levels (normalized to AKT1 levels, right) in SW620-L1 cells treated with EPZ015666 (0, 0.31 or 1.25 μM, for 48 h) and incubated with 200 μM AA (for 15 min) and 800 nM insulin (for 10 min). Three independent western blot analyses were performed. (**E, F)** Representative western blots (E) and quantification of p-mTOR levels (normalized to mTOR levels, F, left), p-AKT levels (normalized to AKT levels, F, middle), and p-4E-BP1 levels (normalized to 4E-BP1 levels, F, right) in HCT-15 cells with PRMT5 overexpression in combination with double knockout of endogenous AKT1 and AKT2 and overexpression of wild-type AKT1 (PRMT5/sgAKT1-1/sgAKT2/AKT1 or PRMT5/sgAKT1-2/sgAKT2/AKT1), or the AKT1 R391K mutant that cannot be methylated by PRMT5 (PRMT5/sgAKT1-1/sgAKT2/AKT1-R391K or PRMT5/sgAKT1-2/sgAKT2/AKT1-R391K) incubated with 200 μM AA (for 15 min) and 800 nM insulin (for 10 min). Three independent western blot analyses were performed. (**G)** *In vitro* proliferation assay of HCT-15 cells with overexpression of PRMT5 and WDR77 in combination with double knockout of endogenous AKT1 and AKT2 and overexpression of wild-type AKT1 (PRMT5/WDR77/sgAKT1-1/sgAKT2/AKT1 or PRMT5/WDR77/sgAKT1-2/sgAKT2/AKT1) or the AKT1-R391K mutant that cannot be methylated by PRMT5 (PRMT5/WDR77/sgAKT1-1/sgAKT2/AKT1 R391K or PRMT5/WDR77/sgAKT1-2/sgAKT2/AKT1 R391K) (1,000 cells) cultured with 200 μM AA and 800 nM insulin for 24 hr. Five independent CCK-8 assays were performed. The data are presented as the mean ± s.e.m. values. *P*-values were determined by unpaired one-way ANOVA with uncorrected Fisher’s LSD test (C and D), or unpaired two-way ANOVA with uncorrected Fisher’s LSD test (F and G). * *P* < 0.05; ** *P* < 0.01; *** *P* < 0.001; **** *P* < 0.0001.

Furthermore, in colorectal cancer cells with double knockout of endogenous AKT1 and AKT2, the ability of PRMT5 to activate the AKT/mTOR signaling pathway in order to increase cell proliferation was restored when AKT1 expression was restored with wild-type AKT1 but not the AKT1 R391K mutant (which contains a mutation blocking its PRMT5-mediated methylation ^[25]^) (Figure 5E-G and Figure S7B-D, Supporting Information). These results suggested that PRMT5 methylated AKT1 at R391 to activate the AKT/mTOR signaling pathway, consequently increasing colorectal cancer cell proliferation. In addition, AKT3 was undetectable in colorectal cancer cells ^[38]^ (Figure S7E, Supporting Information).

### 2.6. NEDD4L promotes PRMT5 degradation to inhibit the AKT/mTOR signaling pathway and prevent colorectal cancer liver metastasis

As demonstrated above, NEDD4L promotes PRMT5 degradation, and PRMT5 methylates AKT1 to activate the AKT/mTOR signaling pathway. Therefore, whether NEDD4L inhibits the AKT/mTOR signaling pathway by promoting PRMT5 degradation to attenuate the arginine methylation of AKT1 was examined. The knockdown of NEDD4L stabilized PRMT5, resulting in arginine methylation of AKT1, whereas the overexpression of NEDD4L-R in NEDD4L-knockdown colorectal cancer cells promoted PRMT5 degradation to attenuate AKT1 methylarginine (Figure 6A). In contrast, the overexpression of wild-type NEDD4L but not the E3 ligase activity-dead NEDD4L C821A mutant promoted PRMT5 degradation to attenuate AKT1 methylarginine (Figure 6B), suggesting that the function of NEDD4L in the arginine methylation of AKT1 is dependent on its E3 ligase activity.

**Figure 6.**
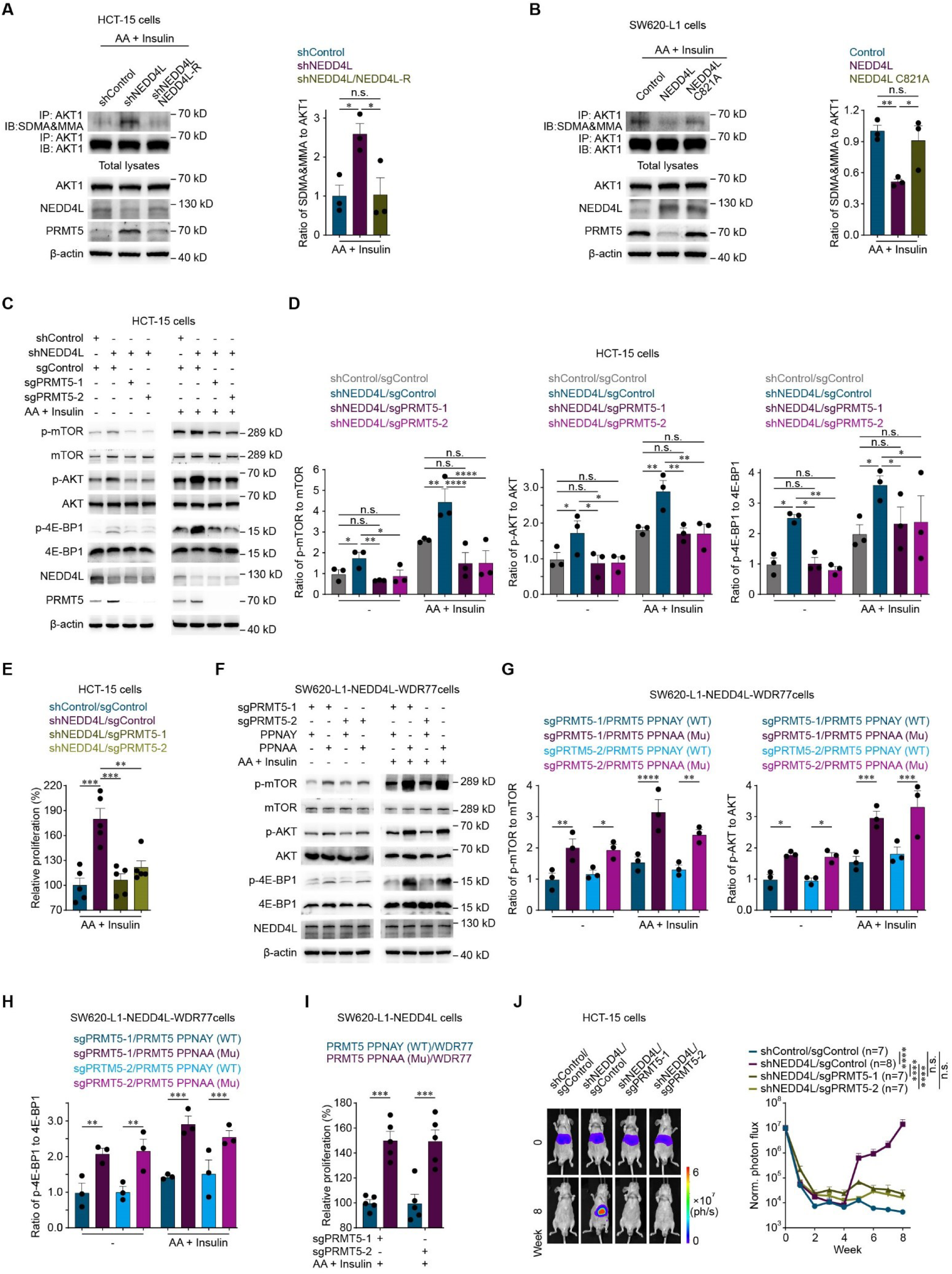
NEDD4L promotes PRMT5 degradation to inhibit the AKT/mTOR signaling pathway and prevent colorectal cancer liver metastasis. (**A)** Representative western blots (left) and quantification of AKT1 methylarginine levels (normalized to AKT1 levels, right) in control HCT-15 cells (shControl), NEDD4L-knockdown HCT-15 cells (shNEDD4L) and HCT-15 cells with knockdown of NEDD4L and restoration with wild-type NEDD4L resistant to shRNA targeting NEDD4L (shNEDD4L/NEDD4L-R) incubated with 200 μM amino acid (AA; for 15 min) and 800 nM insulin (for 10 min). Three independent western blot analyses were performed. (**B)** Representative western blots (left) and quantification of AKT1 methylarginine levels (normalized to AKT1 expression, right) in control SW620-L1 cells (Control) and SW620-L1 cells overexpressing wild-type NEDD4L (NEDD4L) or an E3 ligase activity-dead mutant of NEDD4L (C821A) induced by treatment with 2 µg/ml doxycycline (for 24 hr) and incubated with 200 μM AA (for 15 min) and 800 nM insulin (for 10 min). Three independent western blot analyses were performed. (**C, D)** Representative western blots (C) and quantification of p-mTOR levels (normalized to mTOR levels, D, left), p-AKT levels (normalized to AKT levels, D, middle), and p-4E-BP1 levels (normalized to 4E-BP1 levels, D, right) in control HCT-15 cells (shControl/sgControl), HCT-15 cells with NEDD4L knockdown (shNEDD4L/sgControl), and HCT-15 cells with NEDD4L knockdown in combination with knockout of PRMT5 (shNEDD4L/sgPRMT5-1 or shNEDD4L/sgPRMT5-2) incubated with or without 200 μM AA (for 15 min) and 800 nM insulin (for 10 min). Three independent western blot analyses were performed. (**E)** *In vitro* proliferation assay of control HCT-15 cells (shControl/sgControl), HCT-15 cells with knockdown of NEDD4L (shNEDD4L/sgControl), and HCT-15 cells with knockdown of NEDD4L in combination with knockout of PRMT5 (shNEDD4L/sgPRMT5-1 or shNEDD4L/sgPRMT5-2) (1,000 cells) cultured with 200 μM AA and 800 nM insulin for 24 hr. Five independent CCK-8 assays were performed. (**F-H)** Representative western blots (F) and quantification of p-mTOR levels (normalized to mTOR levels, G, left), p-AKT levels (normalized to AKT levels, G, right), and p-4E-BP1 levels (normalized to 4E-BP1 levels, H) in SW620-L1 cells with overexpression of NEDD4L and WDR77 (NEDD4L-WDR77) in combination with knockout of endogenous PRMT5 (sgPRMT5-1 or sgPRMT5-2) and overexpression of PRMT5 containing the wild-type NEDD4L binding motif (PPNAY) or mutant NEDD4L binding motif (PPNAA) incubated with or without 200 μM AA (for 15 min) and 800 nM insulin (for 10 min). NEDD4L overexpression in cancer cells was induced by treatment with 2 µg/ml doxycycline (for 24 hr). Three independent western blot analyses were performed. (**I)** *In vitro* proliferation assay of SW620-L1 cells with overexpression of NEDD4L and WDR77 (NEDD4L-WDR77) in combination with knockout of endogenous PRMT5 (sgPRMT5-1 or sgPRMT5-2) and overexpression of PRMT5 containing wild-type NEDD4L binding motif (PPNAY) or mutant NEDD4L binding motif (PPNAA) (3,000 cells) cultured with 200 μM AA and 800 nM insulin for 24 hr. Five independent CCK-8 assays were performed. (**J)** Bioluminescence imaging results(left) and quantification of liver metastases (right) in BALB/c nude mice implanted with control HCT-15 cells (shControl/sgControl), HCT-15 cells with knockdown of NEDD4L (shNEDD4L/sgControl), and HCT-15 cells with knockdown of NEDD4L in combination with knockout of PRMT5 (shNEDD4L/sgPRMT5-1 or shNEDD4L/sgPRMT5-2) (3 × 10^6^ cells) via intrasplenic injection. The n-values denote the number of mice per group. The data are presented as the mean ± s.e.m. values. *P*-values were determined by unpaired one-way ANOVA with uncorrected Fisher’s LSD test (A, B, and E), or unpaired two-way ANOVA with uncorrected Fisher’s LSD test (D, G, H, I, and J). * *P* < 0.05; ** *P* < 0.01; *** *P* < 0.001; **** *P* < 0.0001; n.s., not significant.

Moreover, when PRMT5 was knocked out (Figure 6C-E) or when the PRMT5 PPNAA mutant, (whose degradation cannot be mediated by NEDD4L) was overexpressed (Figure 6F-I), NEDD4L failed to inhibit the AKT/mTOR signaling pathway and had no effect on colorectal cancer cell proliferation. Furthermore, the *in vivo* functional experiment revealed that when PRMT5 was knocked out, NEDD4L knockdown failed to promote colorectal cancer liver metastasis (Figure 6J). These results suggested that NEDD4L promoted PRMT5 degradation to inhibit the AKT/mTOR signaling pathway, consequently decreasing colorectal cancer cell proliferation and ultimately preventing colorectal cancer liver metastasis.

Collectively, our findings support the hypothesis that in liver-metastatic colorectal cancer cells, NEDD4L binds to the PPNAY motif in PRMT5, resulting in the ubiquitination and degradation of PRMT5. PRMT5 degradation attenuates the methylation of an arginine residue in AKT1 to inhibit the AKT/mTOR signaling pathway, consequently decreasing colorectal cancer cell proliferation and ultimately preventing colorectal cancer liver metastasis (Figure 7).

**Figure 7.**
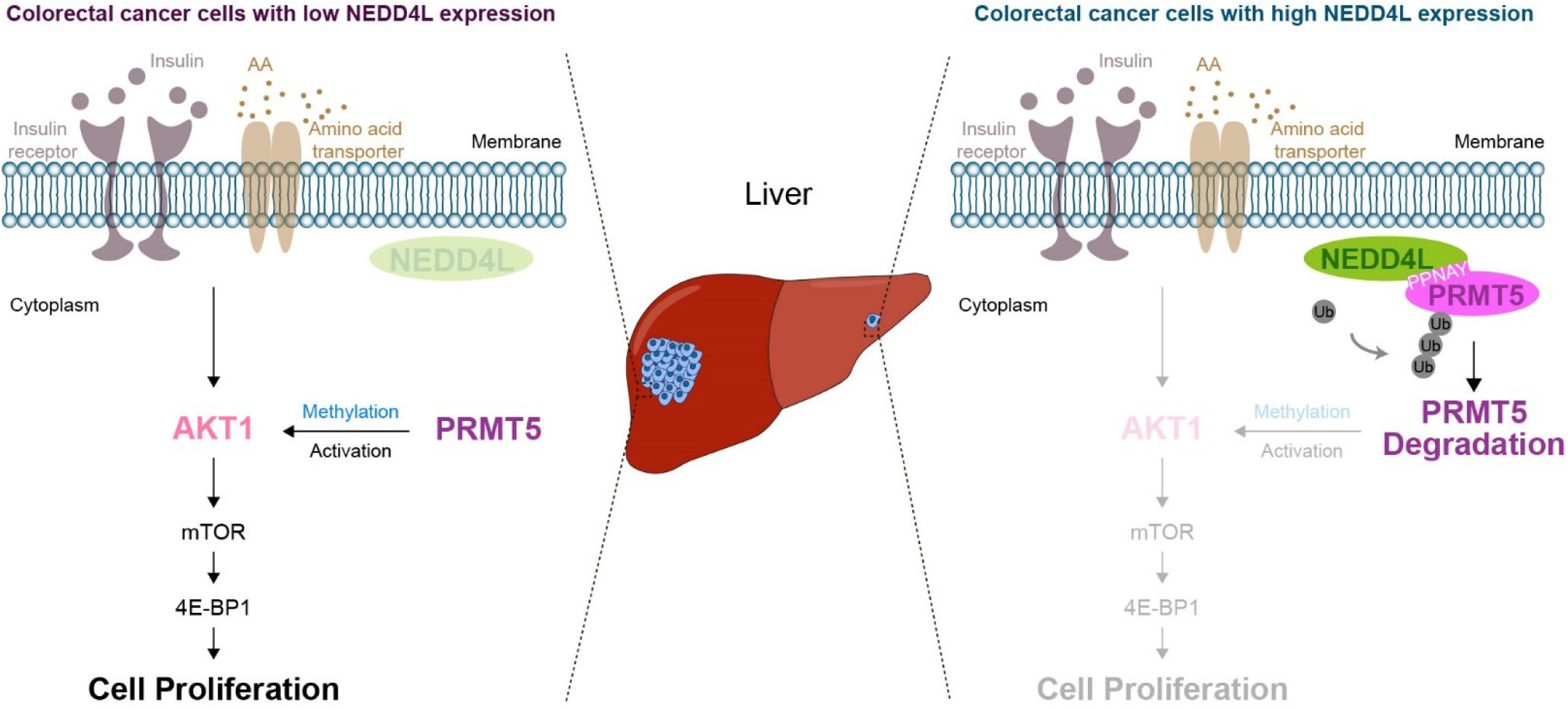
The mechanism by which NEDD4L prevents colorectal cancer liver metastasis. In colorectal cancer cells with high expression of NEDD4L, E3 ligase NEDD4L binds to PPNAY motif in PRMT5 resulting in ubiquitination and degradation of PRMT5. PRMT5 degradation attenuates the methylation of an arginine residue in AKT1 to inhibit AKT/mTOR signaling pathway, consequently decreasing colorectal cancer cell proliferation and ultimately preventing colorectal cancer liver metastasis. Conversely, in colorectal cancer cells with low expression of NEDD4L, the level of NEDD4L is not sufficient for ubiquitination and degradation of PRMT5, resulting in an increase in AKT1 methylarginine to activate the AKT/mTOR signaling pathway. The activation of the AKT/mTOR signaling pathway increases colorectal cancer cell proliferation and ultimately promotes colorectal cancer liver metastasis.

## 3. Discussion

Recently, various reports have shown that dysfunctional activity of E3 ligases is associated with cancer ^[6–8]^. Since E3 ligases play a critical role in the specific recognition of substrate proteins ^[4,5]^, targeting their active sites or interrupting the interactions between E3 ligases and their substrate proteins is a good approach for developing anti-cancer drugs with fewer side effects than currently available drugs ^[39]^. To date, multiple drugs targeting E3 ligases have been approved by the FDA or evaluated in phase I/II clinical trials, for example, thalidomide which suppresses CRBN activity (FDA-approved for the treatment of multiple myeloma), and RG7112, which targets MDM2 to increase the p53 level (evaluated in a phase I clinical trial for hematological malignancies) ^[40]^. Although liver metastasis is a major cause of death in patients with colorectal cancer ^[2]^, there are no drugs that target E3 ligases for the treatment of colorectal cancer liver metastasis. Thus, to identify the key E3 ligase involved in colorectal cancer liver metastasis, we used an shRNA library targeting 156 E3 ubiquitin ligases to perform an *in vivo* loss-of-function screen of a human colorectal cancer cell line in a mouse model of colorectal cancer liver metastasis. Through this screen, five E3 ligases were identified: NEDD4L, BMI1, SMURF1, AREL1 and FBXW2. Among these five candidate E3 ubiquitin ligases, only NEDD4L was identified in two independent liver metastases, and was the top-ranked E3 ligase (Figure S1A, Supporting Information). In colorectal cancer, NEDD4L has been reported to have diverse ubiquitination substrates, such as STK35, YBX1, and Wnt3 ^[15,18,19]^. Previous studies have shown only the suppressive role of NEDD4L and its substrates in primary colorectal cancer ^[12,14,18–20]^. In this study, we focused on metastatic colonization in the liver, which is the rate-limiting step of the metastatic cascade ^[27]^. We revealed that NEDD4L prevents colorectal cancer liver metastasis by inhibiting the proliferation of colorectal cancer cells. In this study, we not only revealed that NEDD4L prevents colorectal cancer liver metastasis but also identified a novel mechanism by which NEDD4L affects colorectal cancer liver metastasis. These results suggest that the E3 ligase NEDD4L could be exploited therapeutically for colorectal cancer liver metastasis.

Notably, we discovered that PRMT5 is a substrate protein of NEDD4L. Accumulating evidence indicates that PRMT5 functions as an oncogene in colorectal cancer and that its cancer-promoting function depends on its methyltransferase activity ^[24–26]^, suggesting that PRMT5 is a potential target for cancer therapy. Indeed, different types of PRMT5 inhibitors have been developed for cancer treatment and are categorized into 5 distinct types: SAM-competitive (JNJ64619178), substrate-competitive (GSK3326595 and EPZ015666), MTA-cooperative (AMG193 and TNG908), allosteric, and PRMT5-adaptor protein-protein interaction inhibitors ^[41]^. Currently, almost all the PRMT5 inhibitors under development have been designed to inhibit PRMT5 methyltransferase activity or disrupt the interactions between the PRMT5 and its target proteins. This study revealed not only that the ubiquitin`‒`proteasome system regulates PRMT5 stability but also that the E3 ligase NEDD4L mediates the ubiquitination and degradation of PRMT5. These findings may provide a new strategy for PRMT5 degradation that differs from the traditional strategies designed to inhibit PRMT5-mediated methylarginine.

The AKT/mTOR signaling pathway is one of the most frequently altered pathways in human cancers ^[42,43]^. Cancer cell proliferation is regulated mainly through AKT-mediated activation of the protein kinase mTORC1 ^[44]^. Both dysfunction of AKT regulators and genetic alterations in AKT cause overactivation of the AKT pathway in various human cancers ^[25]^. Posttranslational modifications of AKT, such as methylarginine, lysine modifications, and tyrosine phosphorylation, are important for AKT hyperactivation in cancers ^[44]^. A recent study revealed that PRMT5 methylates AKT1 at R391 to activate AKT and consequently promote tumorigenesis ^[25]^. In this study, we further demonstrated that PRMT5-mediated R391 methylation of AKT1 activated the AKT/mTOR signaling pathway to increase cell proliferation, which played an important role in colorectal cancer liver metastasis. Taken together, our findings reveal that NEDD4L binds to the PPNAY motif in PRMT5 to induce its ubiquitination and degradation. PRMT5 degradation attenuates the methylation of AKT1 to inhibit the AKT/mTOR signaling pathway, consequently decreasing cancer cell proliferation and ultimately preventing colorectal cancer liver metastasis.

## 4. Conclusions

In this study, we revealed that NEDD4L prevented colorectal cancer liver metastasis, and the metastasis-inhibiting function of NEDD4L was dependent on its E3 ubiquitin ligase activity. We discovered that ubiquitin-proteasome system regulated PRMT5 stability, and PRMT5 was a substrate protein of NEDD4L. Mechanistic studies revealed that NEDD4L bound to the PPNAY motif in PRMT5 and ubiquitinated PRMT5 to promote its degradation. PRMT5 degradation attenuates the methylation of an arginine residue in AKT1. The decrease of methylarginine of AKT1 inhibited the AKT/mTOR signaling pathway, consequently decreasing colorectal cancer cell proliferation. This result may provide a potential target for colorectal cancer liver metastasis therapy.

## 5. Experimental Section

### In vivo loss-of-function screen of E3 ligases involved in colorectal cancer liver metastasis

We selected 794 shRNAs targeting 156 cancer-related E3 ubiquitin ligases (3-9 shRNAs per ligase, Table S1, Supporting Information) to construct an shRNA library for an *in vivo* loss-of-function screen. The sequences of the shRNAs were obtained from Sigma‒Aldrich (USA). The library was subdivided into six subpools. HCT-15 human colorectal cancer cells stably expressing tdTomato-luciferase were transduced independently with each of the six subpools at a multiplicity of infection (MOI) of 3:1, an MOI at which more than 90% of the HCT-15 cells were successfully transduced with the subpools. The transduced HCT-15 cells (3 × 10^6^ cells/100 μl of PBS) were subsequently implanted into BALB/c nude mice via intrasplenic injection (2-3 mice per subpool). Eight weeks after injection, macroscopic metastatic lesions in the liver (visible to the naked eye) were harvested and minced to isolate cancer cells. The clonogenic cancer cells were expanded in medium supplemented with puromycin, and the genomic DNA of the cancer cells was extracted. The resident shRNA-encoding sequences in the genomic DNA were amplified with primers (Table S3, Supporting Information), the PCR products were subsequently cloned and inserted into a TA cloning vector, and at least 15 independent clones were analyzed via Sanger sequencing. Moreover, the mRNA levels of the shRNAs targeting E3 ubiquitin ligases were measured via qPCR to assess the on-target knockdown efficiency of each shRNA.

### Cell lines

The HCT-15, SW480, LS 174T, and COLO 320DM human colorectal cancer cell lines were originally obtained from the American Type Culture Collection (ATCC, USA). The CT26 mice colorectal cancer cell line was originally obtained from ATCC. The SW620-L1 human colorectal cancer cell line was a kind gift from Dr. Filippo G. Giancotti (Columbia University, USA). The 293FT cell line was purchased from Life Technologies. HCT-15 and CT26 cells were cultured in RPMI-1640 medium (31800-089, Thermo Fisher Scientific, USA) supplemented with 10% fetal bovine serum (FBS; A0500-3010, Cegrogen, Germany), 2 mM L-glutamine (21051-024, Thermo Fisher Scientific), and 100 U/ml penicillin/0.1 mg/ml streptomycin (P/S; C0222, Beyotime Biotechnology, China). SW480 and SW620-L1 cells were cultured in DMEM/F12 medium (12500-062, Thermo Fisher Scientific) supplemented with 10% FBS, 2 mM L-glutamine, P/S and 100 μM MEM non-essential amino acids solution (NEAA; 11140050, Thermo Fisher Scientific). LS 174T, COLO 320DM and 293FT cells were cultured in DMEM-HG (12100-061, Thermo Fisher Scientific) supplemented with 10% FBS, 2 mM L-glutamine and P/S. For bioluminescent tracking, cell lines were transduced with a lentiviral vector encoding tdTomato and firefly luciferase, and tdTomato-positive cells were isolated via FACS. The HCT-15, SW620-L1 and CT26 cells used in this study stably expressed both tdTomato and firefly luciferase. All human cell lines were authenticated by short tandem repeat (STR) profiling performed by the Bio-Research Innovation Center Suzhou, SIBCB, CAS, and were routinely tested for mycoplasma contamination.

### Mice

The mice were housed under specific pathogen-free (SPF) conditions in the animal facility of Tongji University. All animal experiments were approved by the Institutional Animal Care and Use Committee of Tongji University (approval number: TJBA01020104). BALB/c nude and BALB/c mice were purchased from the Beijing Vital River Laboratory Animal Technology (Beijing, China). HCT-15 cells, SW620-L1 cells, and their derivatives were used to establish xenograft models in 5- to 7-week-old male BALB/c nude mice. CT-26 cells, and their derivatives were used to establish isograft models in 5- to 7-week-old male syngeneic BALB/c mice.

### Bioluminescence imaging

Mice were anesthetized and injected retro-orbitally with 1.5 mg of D-luciferin (LUCK-1G, Gold Biotechnology, USA) at the indicated times.

The animals were imaged in a NightOWL II LB 983 chamber (BertholdTechnologies, Bad Wildbad, Germany), AniView 100 (Biolight Biotechnology, China) or IVIS Lumina XRMS Series III (PerkinElmer, USA) within 5 min after D-luciferin injection, and data were recorded with Indigo™ software (BertholdTechnologies, Bad Wildbad, Germany), AniView software (Biolight Biotechnology, China) or Live Image software (PerkinElmer, USA) to measure liver metastasis. Photon flux was calculated in a circular region of interest (ROI) encompassing the liver of each mouse.

### Establishment of mouse models of colorectal cancer liver metastasis via intrasplenic injection

Cancer cells stably expressing tdTomato and firefly luciferase were harvested by trypsinization, washed twice in PBS, and resuspended at 3 × 10^6^ (transduced shControl/Control, shNEDD4L/Control, shNEDD4L/NEDD4L-R, Control, PRMT5/WDR77 or PRMT5 R368A/WDR77 HCT-15 cells), 3 × 10^6^ (transduced shControl/sgControl, shNEDD4L/sgControl, shNEDD4L/sgPRMT5-1 or shNEDD4L/sgPRMT5-2 HCT-15-Cas9 cells), 1 × 10^6^ (transduced inducible Control, NEDD4L, NEDD4L C821A or NEDD4L R776Q SW620-L1 cells), 1 × 10^6^ (transduced sgControl, sgPRMT5-1 or sgPRMT5-2 SW620-L1-Cas9 cells), or 3 × 10^5^ (transduced induced Control or NEDD4L CT26 cells) cells in 100 μl of PBS were injected into the spleens of the mice. The spleens were excised 2 min after cancer cell injection. Bioluminescence imaging was used to verify successful injection after surgery and to monitor metastatic outgrowth. To induce the overexpression of NEDD4L or its mutants in SW620-L1 cells, doxycycline (D9891, Sigma; 100 μg doxycycline in 100 μl of PBS) was administered intraperitoneally on the day of cancer cell injection (day 0) and was then administered orally (400 ppm in chow combined with 2 mg/ml in water) from day 0 to the experimental endpoint. To inhibit the mTOR signaling pathway *in vivo*, rapamycin (GC15031, GlpBio, USA; 2 mg/kg) was administered intraperitoneally every two days from the day of cancer cell injection (day 0) to the experimental endpoint.

### Immunostaining

Cancer cells stably expressing tdTomato and firefly luciferase were harvested by trypsinization, washed twice in PBS, resuspended (1 × 10^6^ transduced inducible Control or NEDD4L SW620-L1 cells in 100 μl of PBS), and injected into the spleens of mice. The spleens were excised 2 min after cancer cell injection. To induce NEDD4L overexpression in SW620-L1 cells, doxycycline (100 μg doxycycline in 100 μl of PBS) was administered intraperitoneally on day 7 after cancer cell injection and was then administered orally (400 ppm in chow combined with 2 mg/ml in water) from day 7 to day 13 after cancer cell injection. Mice were sacrificed and perfused with PBS through the inferior vena cava on day 13 after cancer cell injection. The entire liver was divided into 7 parts based on the liver lobes. The liver lobes were fixed with 4% paraformaldehyde (PFA) overnight at 4°C, dehydrated in 30% sucrose for 48 h, and embedded in optimal cutting temperature (OCT) compound. The entire liver lobe was serially sectioned (10 μm thick) with a Thermo Scientific Cryostar NX50, and immunofluorescence staining was performed. An anti-tdTomato antibody was used with a Tyramide Signal Amplification Kit (B40931, Thermo Fisher Scientific) to visualize cancer cells, and staining with an anti-Ki67 antibody was then performed to assess proliferation. Immunoreactions were detected with fluorescently labeled secondary antibodies (Thermo Fisher Scientific). The sections were mounted with ProLong™ Gold Antifade Mountant with DAPI (P36931, Thermo Fisher Scientific) and imaged with a Carl Zeiss LSM900 confocal microscope. The metastatic lesions in the entire liver were counted at 40× magnification. For each metastatic lesion, three random sections were selected, the Ki67-positive rate was determined by calculating the percentage of Ki67-positive cells among the total population of tumor cells in each section, and the percentage of Ki67-positive cells in each metastatic lesion was then calculated as the average percentage across the three sections. Since there were no metastatic lesions or cells in the liver in the NEDD4L group, the percentage of Ki67-positive cells was considered 0.

### CCK-8 assay

HCT-15 cells transduced with shControl/Control, shNEDD4L/Control, or shNEDD4L/NEDD4L-R (1,000 cells) were seeded into 96-well plates and cultured for 72 hours. SW620-L1 cells transduced with Control, NEDD4L, NEDD4L C821A, or NEDD4L R776Q (3,000 cells) were seeded into seeded into 96-well plates and cultured with 2 μg/ml doxycycline for 72 hours. At the indicated time points, CCK-8 solution (CK04, Dojindo, Japan) was added to each well, and the plates were incubated for 1 hour at 37°C. The absorbance was measured at 450 nm with a spectrophotometer (Infinite M200 Pro, TECAN, Switzerland). HCT-15 cells transduced with shControl/Control, shNEDD4L/Control, shNEDD4L/NEDD4L-R, Control, PRMT5/WDR77, or PRMT5 R368A/WDR77 (1,000 cells), HCT-15-Cas9 cells transduced with shControl/sgControl, shNEDD4L/sgControl, shNEDD4L/sgPRMT5-1, shNEDD4L/sgPRMT5-2, sgAKT1-1/sgAKT2/AKT1/PRMT5/WDR77, sgAKT1-1/sgAKT2/AKT1 R391K/PRMT5/ WDR77, sgAKT1-2/sgAKT2/AKT1/PRMT5/WDR77, or sgAKT1-2/sgAKT2/AKT1 R391K/PRMT5/WDR77 (1,000 cells), SW620-L1 cells transduced with Control, NEDD4L, NEDD4L C821A, or NEDD4LR776Q (3,000 cells), or SW620-L1-Cas9 cells transduced with sgControl, sgPRMT5-1, sgPRMT5-2, NEDD4L/sgPRMT5-1/PRMT5-PPNAY, NEDD4L/sgPRMT5-1/PRMT-PPNAA, NEDD4L/sgPRMT5-2/PRMT5-PPNAY, or sgPRMT5-2/NEDD4L/PRMT5-PPNAA (3,000 cells) were seeded into 96-well plates and cultured for 48 hours. The medium was replaced with FBS-free RPMI-1640 medium (for HCT-15 cells, 31800-089, Thermo Fisher Scientific) or DMEM/F12 medium (for SW620-L1 cells, 12500-062, Thermo Fisher Scientific), and the cancer cells were cultured for another 24 hours. The medium was subsequently replaced with FBS-free, amino acid-free RPMI-1640 medium (for HCT-15 cells, YYX2009, Yuyan Biotechnology, China) or DMEM/F12 medium (for SW620-L1 cells, YYX2010, Yuyan Biotechnology) supplemented with 800 nM insulin (I9278, Sigma) and 200 μM nonessential amino acids (NEAA; 11140050, Thermo Fisher Scientific), and the cancer cells were cultured for another24 hours. To induce the overexpression of NEDD4L or its mutants in SW620-L1 cells, cells were cultured with 2 µg/ml doxycycline. To inhibit the mTOR signaling pathway in HCT-15 cells transduced with shControl, shNEDD4L, Control, PRMT5/WDR77 or PRMT5-R368A/WDR77, cells were cultured with 100 nM rapamycin for 24 hours. At the indicated time points, CCK-8 solution was added to each well, and the plates were incubated for 2 hours at 37°C. The absorbance was measured at 450 nm with a spectrophotometer (Infinite M200 Pro, TECAN).

### Coimmunoprecipitation (Co-IP) assay

To confirm the interaction between endogenous PRMT5 and AKT1, SW620-L1 cells were seeded in a 10 cm dish and cultured for 48 hours. The medium was replaced with FBS-free DMEM/F12, and the cells were cultured for 24 hours. The medium was then replaced with FBS-free, amino acid-free DMEM/F12, and the cells were cultured for 50 min prior to incubation with 200 μM NEAA for 15 min and 800 nM insulin for 10 min. After treatment, the cells were lysed in co-IP radioimmunoprecipitation assay (RIPA) lysis buffer (50 mM Tris-HCl (pH 7.4), 150 mM NaCl, 1% NP-40, and 0.25% sodium deoxycholate) supplemented with phenylmethylsulfonyl fluotide (PMSF; ST505, Beyotime Biotechnology) and phosphatase inhibitor cocktail (B15001, Bimake, USA). To immunoprecipitate PRMT5, total lysates containing 1 mg of protein were incubated with 1 µg of a rabbit polyclonal anti-PRMT5 antibody overnight at 4°C and were then incubated with 20 µl of Protein A/G agarose (22851, Thermo Fisher Scientific) for 3 hours at 4°C. The immunoprecipitates and total lysates were then subjected to immunoblotting with the indicated antibodies (Table S4, Supporting Information). To confirm the interactions between exogenous Flag-PRMT5 and AKT/mTOR signaling pathway-related proteins (HA-AKT, HA-TSC2 and Myc-Rheb), 293FT cells transfected with control or Flag-PRMT5/Myc-WDR77 in combination with HA-AKT, HA-TSC2 or Myc-Rheb were cultured for 48 hours. The medium was then replaced with FBS-free DMEM-HG (12100-061, Thermo Fisher Scientific), and the cells were cultured for another 24 hours. Then, the medium was replaced with FBS-free, amino acid-free DMEM (CB000-8001, Excell Bio, China), and the cells were cultured for 50 min prior to incubation with 200 μM NEAA for 15 min and 800 nM insulin for 10 min. After treatment, the cells were lysed in co-IP RIPA lysis buffer (50 mM Tris-HCl (pH 7.4), 150 mM NaCl, 1% NP-40, and 0.25% sodium deoxycholate) supplemented with PMSF (ST505, Beyotime Biotechnology) and phosphatase inhibitor cocktail (B15001, Bimake). To immunoprecipitate Flag-PRMT5, total lysates containing 1 mg of protein were incubated with 20 µl of Anti-FLAG M2 Affinity Gel (A2220, Sigma) for 2 hours at 4°C. The immunoprecipitates and total lysates were then subjected to immunoblotting with the indicated antibodies (Table S3, Supporting Information).

### Mass spectrometry

To identify the substrate of NEDD4L, 293FT cells were transiently transfected with the control plasmid or with Flag-NEDD4L and HA-ubiquitin. After 48 h, the cells were treated with 20 μM MG132 (GC10383, GlpBio) for 12 h and were then lysed in lysis buffer (50 mM Tris-HCl (pH 7.4), 150 mM NaCl, 1 mM EDTA, 1% glycerol, 1% Triton X-100, and 0.1% sodium dodecyl sulfate (SDS)) supplemented with 1 mM Na3VO4, phosphatase inhibitor cocktail (B15001, Bimake), and protease inhibitors (539134, Merck, Germany). The lysates were then incubated with Anti-FLAG M2 Affinity Gel (A2220, Sigma) for 2 hours at 4°C. After washing three times, the protein-gel mixture was eluted with Flag peptide (A6001, Apexbio, USA) for 30 min according to the manufacturer’s protocols. For serial co-IP, the immunoprecipitates obtained with Anti-FLAG M2 Affinity Gel were incubated with EZview^TM^ Red Anti-HA Affinity Gel (E6779, Sigma) for 12 hours at 4°C. After washing three times, the protein-gel mixture was subjected to elution with HA peptide (A6004, Apexbio) for 30 min according to the manufacturer’s protocols. The immunoprecipitates were separated on a NuPAGE 10% Bis-Tris Gel (NP0341BOX, Thermo Fisher Scientific) and visualized with EZBlue™ Gel Staining Reagent (G1041-500ML, Sigma). The complete gel lanes were excised and subjected to electrospray ionization-liquid chromatography-tandem mass spectrometry (ESI-LC-MS/MS) analysis.

### Analysis of protein expression

Cancer cells were lysed in RIPA buffer (50 mM Tris-HCl (pH 7.4), 150 mM NaCl, 1 mM EDTA, 1% Triton X-100, 1% sodium deoxycholate, and 0.1% SDS) supplemented with PMSF and phosphatase inhibitor cocktail unless otherwise noted. Protein concentrations were measured with an Enhanced BCA Protein Assay Kit (P0010, Beyotime). The total lysates were subjected to immunoblotting with the indicated antibodies (Table S4, Supporting Information). Protein expression was quantified with ImageJ.

### Analysis of protein methylarginine levels

SW620-L1-Cas9 cells transduced with sgControl, sgPRMT5-1, or sgPRMT5-2, SW620-L1 cells transduced with Control, NEDD4L, or NEDD4L C821A, HCT-15-Cas9 cells transduced with sgAKT1-1/AKT1/PRMT5/WDR77, sgAKT1-1/AKT1 R391K/PRMT5/WDR77, sgAKT1-2/AKT1/PRMT5/WDR77, or sgAKT1-2/AKT1 R391K/PRMT5/WDR77, HCT-15 cells transduced with shControl/Control, shNEDD4L/Control, or shNEDD4L/NEDD4L-R and HCT-15 cells transduced with Control, PRMT5/WDR77, or PRMT5 R368A/WDR77 (5 × 10^5^ cells) were seeded in a 10 cm dish and cultured for 48 hours. The medium was replaced with FBS-free DMEM/F12 (for SW620-L1 cells) or RPMI-1640 (for HCT-15 cells) and the cells were cultured for 24 hours. The medium was then replaced with FBS-free, amino acid-free DMEM/F12 (for SW620-L1 cells) or RPMI-1640 (for HCT-15 cells), and the cells were cultured for 50 min prior to incubation with 200 μM NEAA for 15 min and 800 nM insulin for 10 min. To induce the overexpression of NEDD4L or NEDD4L C821A in SW620-L1 cells, cells were cultured with 2 µg/ml doxycycline. SW620-L1 cells (5 × 10^5^ cells) were seeded in a 10 cm dish and cultured for 48 hours. Then, the medium was replaced with FBS-free DMEM/F12, and the cells were cultured with EPZ015666 (0, 0.31 or 1.25 μM) for 48 hours. The medium was then replaced with FBS-free, amino acid-free DMEM/F12 for 50 min prior to incubation with 200 μM NEAA for 15 min and 800 nM insulin for 10 min. After treatment, the cells were lysed in RIPA lysis buffer supplemented with PMSF and phosphatase inhibitor cocktail. To analyze the AKT1 methylarginine level, AKT1 was immunoprecipitated. Total lysates containing 1 mg of protein were incubated with 1 µg of a rabbit monoclonal anti-AKT1 antibody overnight at 4°C and were then incubated with 20 µl of Protein A/G agarose for 3 hours at 4°C. To analyze the pan-AKT methylarginine level, pan-AKT was immunoprecipitated. Total lysates containing 1 mg of protein were incubated with 1 µg of a rabbit monoclonal antibody anti-AKT overnight at 4°C and were then incubated with 20 µl of Protein A/G agarose for 3 hours at 4°C. Immunoprecipitates and total lysates were subjected to immunoblotting with the indicated antibodies (Table S4, Supporting Information).

### Analysis of protein ubiquitination levels

SW620-L1 cells transduced with Control, NEDD4L, NEDD4L C821A or NEDD4L R776Q (5 × 10^5^ cells) were seeded into 10 cm dishes and cultured for 48 hours. Subsequently, 20 μM MG132 was added, and the cells were incubated for 12 hours. After treatment, the cells were lysed in co-IP RIPA lysis buffer supplemented with PMSF and phosphatase inhibitors. To induce the overexpression of NEDD4L or its mutants in SW620-L1 cells, cells were cultured with 2 µg/ml doxycycline. To immunoprecipitate PRMT5, total lysates containing 1 mg of protein were incubated with 1 μg of a rabbit monoclonal anti-PRMT5 antibody overnight at 4°C and were then incubated with 20 μl of Protein A/G Plus Agarose for 3 hours at 4°C. 293FT cells transfected with Control/HA-Ubiquitin, NEDD4L/HA-Ubiquitin, NEDD4L C821A/HA-Ubiquitin or NEDD4L R776Q/HA-Ubiquitin (1 × 10^6^ cells) were seeded into 10 cm dishes and cultured for 48 hours. Subsequently, 20 μM MG132 was added, and the cells were incubated for 12 hours. After treatment, the cells were lysed in co-IP RIPA lysis buffer supplemented with PMSF and phosphatase inhibitors. To immunoprecipitate Flag-NEDD4L, total lysates containing 1 mg of protein were incubated with 20 μl of Anti-FLAG M2 Affinity Gel for 2 hours at 4°C. Immunoprecipitates and total lysates were subjected to immunoblotting with the indicated antibodies (Table S4, Supporting Information).

### Antibodies

All the antibodies and their application were listed in Table S4, Supporting Information.

### Analysis of mRNA expression

Total RNA was extracted with an RNAprep Pure Cell/Bacteria Kit (DP430, TIANGEN, China) and reverse transcribed with a ReverTra Ace qPCR RT Kit (FSQ-101, TOYOBO, Japan). An amount of cDNA corresponding to approximately 10 ng of input RNA was used for each reaction. qPCR was performed with a TaqMan Gene Expression Assay (Applied Biosystems, USA) or 2 × SYBR Green qPCR Master Mix (Low ROX, B21702, Bimake). All quantities were normalized to those of endogenous *ACTB* (β-actin). All the experiments were performed on an Applied Biosystems 7500/7500 Fast instrument. The specific TaqMan gene expression assays and the sequences of the primers used to amplify the genes are listed in Table S3, Supporting Information.

### Plasmids

The constructs encoding the shRNAs against human NEDD4L (TRCN0000000905) and human PRMT5 (#1, TRCN0000107088 and #2, TRCN0000107086) were generated by cloning the corresponding shRNA sequences into the pLKO.1 vector. The cDNAs encoding full-length human NEDD4L, PRMT5, WDR77, AKT, AKT1 and Ubiquitin were cloned and sequenced. The pCDNA3.1-HA-TSC2 and pCDNA3.1-Myc-Rheb plasmids were kind gifts from Dr. Ping Wang (Tongji University, China). Flag-NEDD4L, Flag-NEDD4L C821A, and Flag-NEDD4L R776Q were subcloned and inserted into the pCW lentiviral vector or pQCXIP (Clontech, Japan) retroviral vector and verified by sequencing. HA-AKT, HA-ubiquitin, Flag-PRMT5, Myc-WDR77, Myc-PRMT5-PPNAY, Myc-PRMT5-PANAY, Myc-PRMT5-PPNAA, Myc-PRMT5-PPNYY, Flag-PRMT5-2A-Myc-WDR77 and Flag-PRMT5/R368A-2A-Myc-WDR77 were subcloned and inserted into the pQCXIP (Clontech) retroviral vector and verified by sequencing. Flag-NEDD4L-R, Myc-PRMT5, Myc-PRMT5-PPNAA, Flag-PRMT5-2A-Myc-WDR77 were subcloned and inserted into the pQCXIN (Clontech) retroviral vector and verified by sequencing. Flag-AKT1 and Flag-AKT1/R391K were subcloned and inserted into the pBABE-Puro (Clontech) retroviral vector and verified by sequencing. To construct the single guide RNA (sgRNA) expression vector, the sgPRMT5-1 and sgPRMT5-2 sequences were cloned and inserted into the lentiGuide-Puro lentiviral vector or lentiGuide-EGFP lentiviral vector, the sgAKT1-1 and sgAKT1-2 sequences were cloned and inserted into the lentiGuide-EGFP lentiviral vector, and the sgAKT2 sequence was cloned and inserted into the lentiGuide-BFP vector. All guide sequences are listed in Table S5, Supporting Information. Populations of overexpression, knockdown, or knockout cells were used, and the overexpression, knockdown, or knockout efficiency was verified via western blotting.

### Matrigel invasion assay

HCT-15 cells transduced with shControl or shNEDD4L (1,000 cells) were cultured in RPMI-1640 medium on the Matrigel-coated (356237, Corning, USA; 100 μg/well) membranes of transwell cell culture inserts. After a 6-h incubation in RPMI-1640 supplemented with 10% FBS, the remaining cells and Matrigel in the upper compartment of the inserts were removed by wiping the upper side of the membrane with cotton swabs. The invaded cells in the inserts were subsequently fixed with 4% PFA for 10 min at room temperature and stained with crystal violet (C0121, Beyotime). All invaded cells in each insert were counted at 400× magnification (3 inserts per group).

### Tumor sphere formation assay

HCT-15 cells transduced with shControl or shNEDD4L (1,000 cells) were seeded in 24-well ultralow-attachment plates (Corning) and cultured for 7 days in serum-free MEGM (Lonza, Switzerland) supplemented with B27 (1:50; Thermo Fisher Scientific), 20 ng/ml EGF (Thermo Fisher Scientific), 10 ng/ml bFGF (Thermo Fisher Scientific), and 2 μg/ml heparin (Sigma). All tumor spheres in each well were counted.

### Survival analysis

The colorectal cancer dataset GSE17536, the signal intensity matrices calculated with MAS5 and clinical information were downloaded from the Gene Expression Omnibus (GEO) website. The gene expression values for each patient were log2 scaled and converted to z scores. The best probe to represent NEDD4L was selected with the R package “JetSet”. The RNA-seq data matrices and clinical information of the patients represented in the TCGA colorectal cancer (COAD+READ) datasets were downloaded from cBioPortal (https://www.cbioportal.org/datasets). The gene expression values for each patient were log2 scaled and converted to z scores. Clinical information for each patient with an available survival time was collected for survival analysis. The statistical significance of differences in survival was evaluated by Kaplan-Meier analysis with the log-rank test (implemented with the “survdiff” function in the “survival” package in R). The data were plotted as Kaplan-Meier survival curves.

### GSEA

GSEA was performed with the GSEA platform (GSEA v4.0.3). The proliferation-related pathways were analyzed. The patients were classified by their NEDD4L expression level (high- and low-expression groups) as used for survival analysis. The default values were used, except that the metric for ranking genes was set to “t test” and the permutation type was set to “gene set”.

### Analysis of NEDD4L mRNA expression in primary tumors and liver metastatic lesions of patients with colorectal cancer

Five publicly available microarray colorectal cancer datasets (GSE10961, GSE18462, GSE28702, GSE40367, and GSE41568) in the GPL570 platform (Affymetrix Human Genome U133 Plus 2.0 Array) were downloaded from the GEO website. Gene expression was summarized by normalizing the data in each raw CEL file with the MAS5 algorithm in the R statistical environment (www.r-project.org) with the Affy Bioconductor library. The primary tumor and liver metastatic lesions of the patients with colorectal cancer were unpaired. A publicly available microarray colorectal cancer dataset (GSE14297) was downloaded from the GEO website. In this dataset, the primary tumor and liver metastatic lesions of the patients with colorectal cancer were paired.

### Statistical analysis

The group sizes used for the *in vivo* and *in vitro* experiments were selected on the basis of intragroup variation. Data analyses and figure plotting were performed with GraphPad Prism (version 8.3.0). The numbers of replicates and independent experiments are listed in the figure legends. The values are presented as the mean ± s.e.m. values unless otherwise stated. For all analyses, a *P*-value of < 0.05 was considered to indicate statistical significance.

## Supporting Information

Supporting Information is available from the Wiley Online Library or from the author.

## Supporting information

NEDD4L-Supporting Information

## Acknowledgements

We thank members of the Gao laboratory for discussions. This work was supported by the National Natural Science Foundation of China (82303416 and 81972736), and the startup funding from Shanghai Tenth People’s Hospital.

## Author contributions

Z.D., X.S., J.M., Q.C., and Y.G. designed, performed, and analyzed most experiments with assistance from R.C. X.S. performed the bioinformatic analyses. X.S. and H.G. wrote the manuscript with input from Z.D. H.Q. supported the study. H.G. conceived, designed, interpreted, and supervised the study.

## Data Availability Statement

No datasets and materials were generated during the current study.

