## Supplementary material for "The E3 ligase NEDD4L prevents colorectal cancer liver metastasis via degradation of PRMT5 to inhibit the AKT/mTOR signaling pathway": NEDD4L-Supporting Information

**This PDF file includes:**

Figures S1 to S7

Tables S1 to S5

**A**

Results of *in vivo* functional shRNA library targeting human E3 ubiquitin ligases screen.

| Gene ID | Target Genes | No. of mets | shRNA ID |
|---|---|---|---|
| **23327** | **NEDD4L: NEDD4 like E3 ubiquitin protein ligase** | **2 mets from 2 mice** | **TRCN0000000905** |
| 648 | BMI1: B lymphoma Mo-MLV insertion region | 1 mets | TRCN0000020158 |
| 57154 | SMURF1: SMAD specific E3 ubiquitin protein ligase 1 | 1 mets | TRCN0000003472 |
| 9870 | AREL1: apoptosis resistant E3 ubiquitin protein ligase 1 | 1 mets | TRCN0000004425 |
| 26190 | FBXW2: F-box and WD repeat domain containing 2 | 1 mets | TRCN0000006549 |

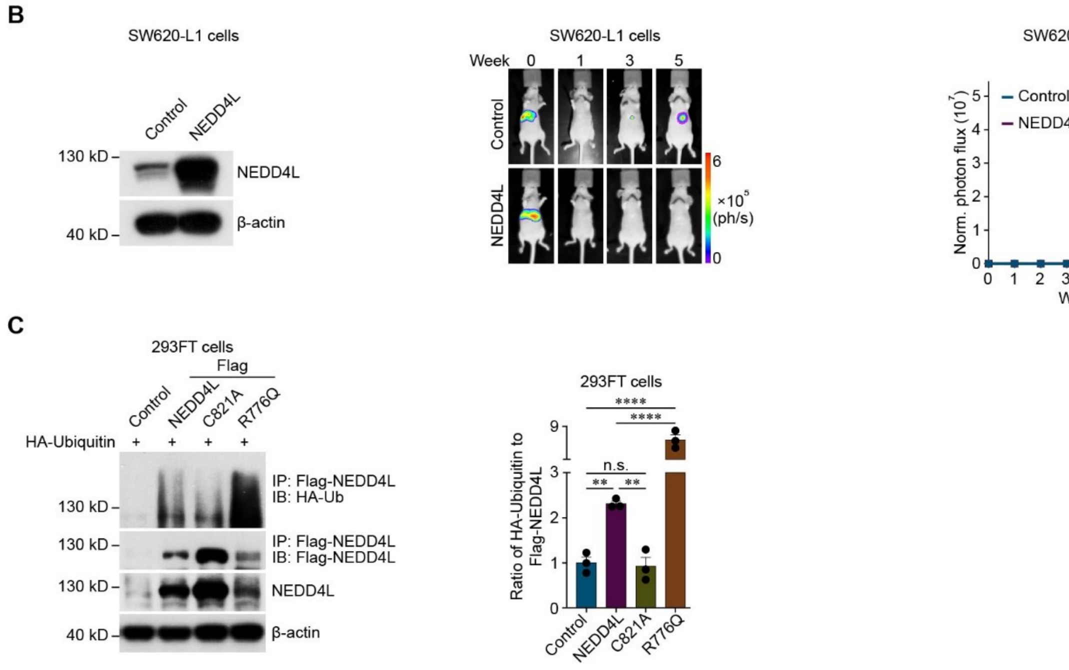


**Figure S1**

**Figure S1.** NEDD4L prevents colorectal cancer liver metastasis.

**(A)** Candidate human E3 ligases identified via the *in vivo* functional screen.

**(B)** Representative western blots showing NEDD4L expression in control SW620-L1 cells (Control) and SW620-L1 cells overexpressing wild-type NEDD4L (NEDD4L) (left). Bioluminescence imaging results (middle) and quantification of liver metastases (right) in BALB/c nude mice implanted with Control or NEDD4L SW620-L1 cells ($1 \times 10^6$ cells) via intrasplenic injection. To induce the overexpression of NEDD4L in the cells, doxycycline (100 μg/mouse) was administered intraperitoneally on the day of cancer cells injection (day 0) and was then administered orally (400 ppm in chow combined with 2 mg/ml in water) from day 0 to the experimental endpoint. The nvalues denote the number of mice per group, and three independent western blot analyses were performed.

**(C)** Representative western blots (left) and qualification of Flag-NEDD4L ubiquitination (normalized to Flag-NEDD4L or its mutant expression, right) in control 293FT cells (Control) and 293FT cells overexpressing wild-type NEDD4L (NEDD4L), an E3 ligase activity-dead mutant of NEDD4L (C821A) or a constitutively active mutant of NEDD4L (R776Q). Three independent western blot analyses were performed.

The data are presented as the mean ± s.e.m. values. *P*- values were determined by unpaired two-way ANOVA with uncorrected Fisher's LSD test (B), or unpaired one-way ANOVA with uncorrected Fisher's LSD test (C). ** $P < 0.01$; *** $P < 0.001$; **** $P < 0.0001$; n.s., not significant.

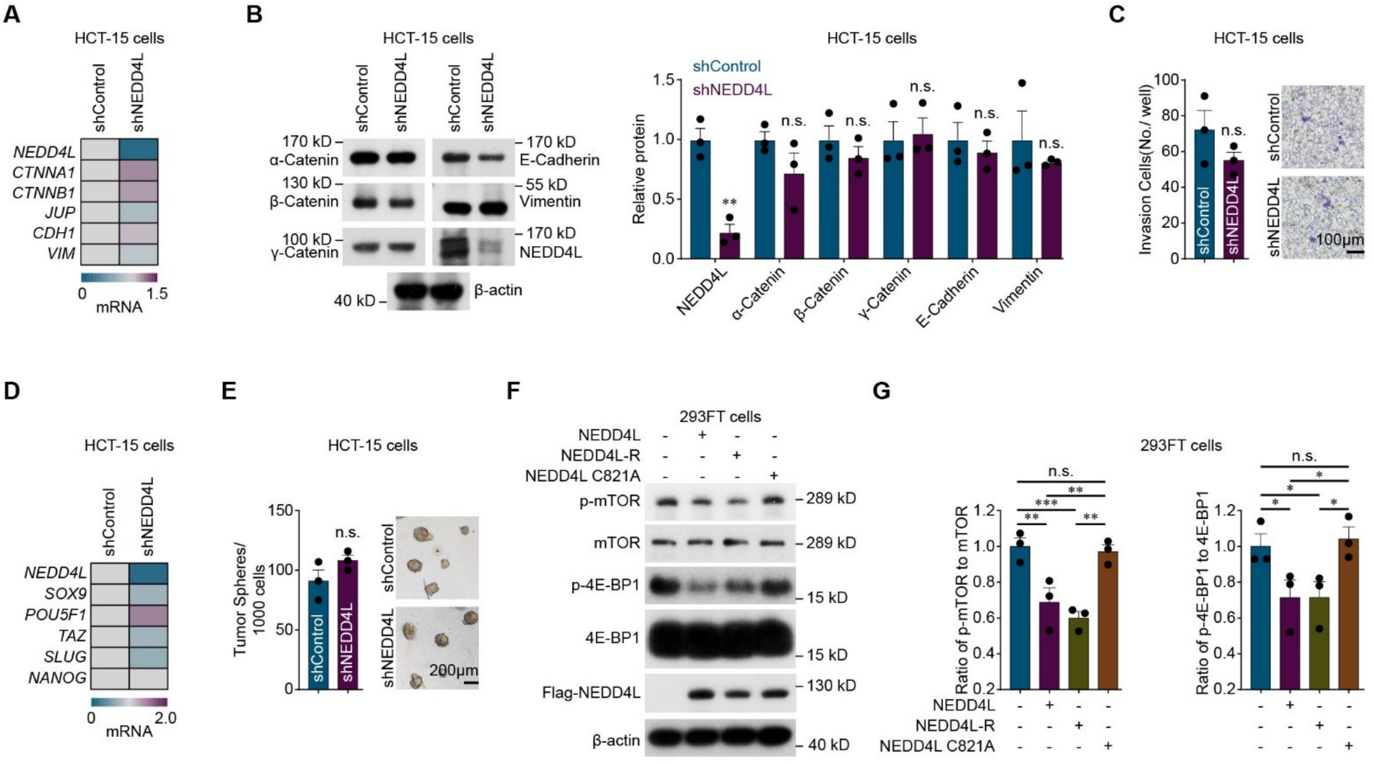


**Figure S2**

**Figure S2.** NEDD4L decreases colorectal cancer cell proliferation by inhibiting the mTOR signaling pathway.

**(A)** qPCR analysis of the mRNA expression of EMT markers in control (shControl) and NEDD4L-knockdown (shNEDD4L) HCT-15 cells. Three independent experiments were performed.

**(B)** Representative western blots (left) and quantification of the protein expression of EMT markers (normalized to β-actin expression, right) in control (shControl) and NEDD4L-knockdown (shNEDD4L) HCT-15 cells. Three independent western blot analyses were performed.

**(C)** Quantification (left) and representative images (right) of the Matrigel invasion ability of control (shControl) and NEDD4L-knockdown (shNEDD4L) HCT-15 cells (1,000 cells) Three independent assays were performed. Scale bar, 100 μm.

**(D)** qPCR analysis of the mRNA expression of stem cell transcription factors in control (shControl) and NEDD4L-knockdown (shNEDD4L) HCT-15 cells. Three independent experiments were performed.

**(E)** Quantification (left) and representative images (right) of tumor sphere assay of control (shControl) and NEDD4L-knockdown (shNEDD4L) HCT-15 cells (1,000 cells). Three independent assays were performed. Scale bar, 200 μm.

**(F, G)** Representative western blots (F) and quantification of p-mTOR levels (normalized to mTOR levels, G, left), and p-4E-BP1 levels (normalized to 4E-BP1levels, G, right) in control 293FT cells (Control) and 293FT cells with overexpression of wild-type NEDD4L (NEDD4L), wild-type NEDD4L resistant to shRNA targeting NEDD4L (NEDD4L-R) or an E3 ligase activity-dead mutant of NEDD4L (C821A). Three independent western blot analyses were performed.

The data are presented as the mean ± s.e.m. values. *P*- values were determined by unpaired two-way ANOVA with uncorrected Fisher's LSD test (B), unpaired two-tailed Student's t-test with Welch's correction (C and E), or unpaired one-way ANOVA with uncorrected Fisher's LSD test (G). * $P < 0.05$; ** $P < 0.01$; *** $P < 0.001$; n.s., not significant.

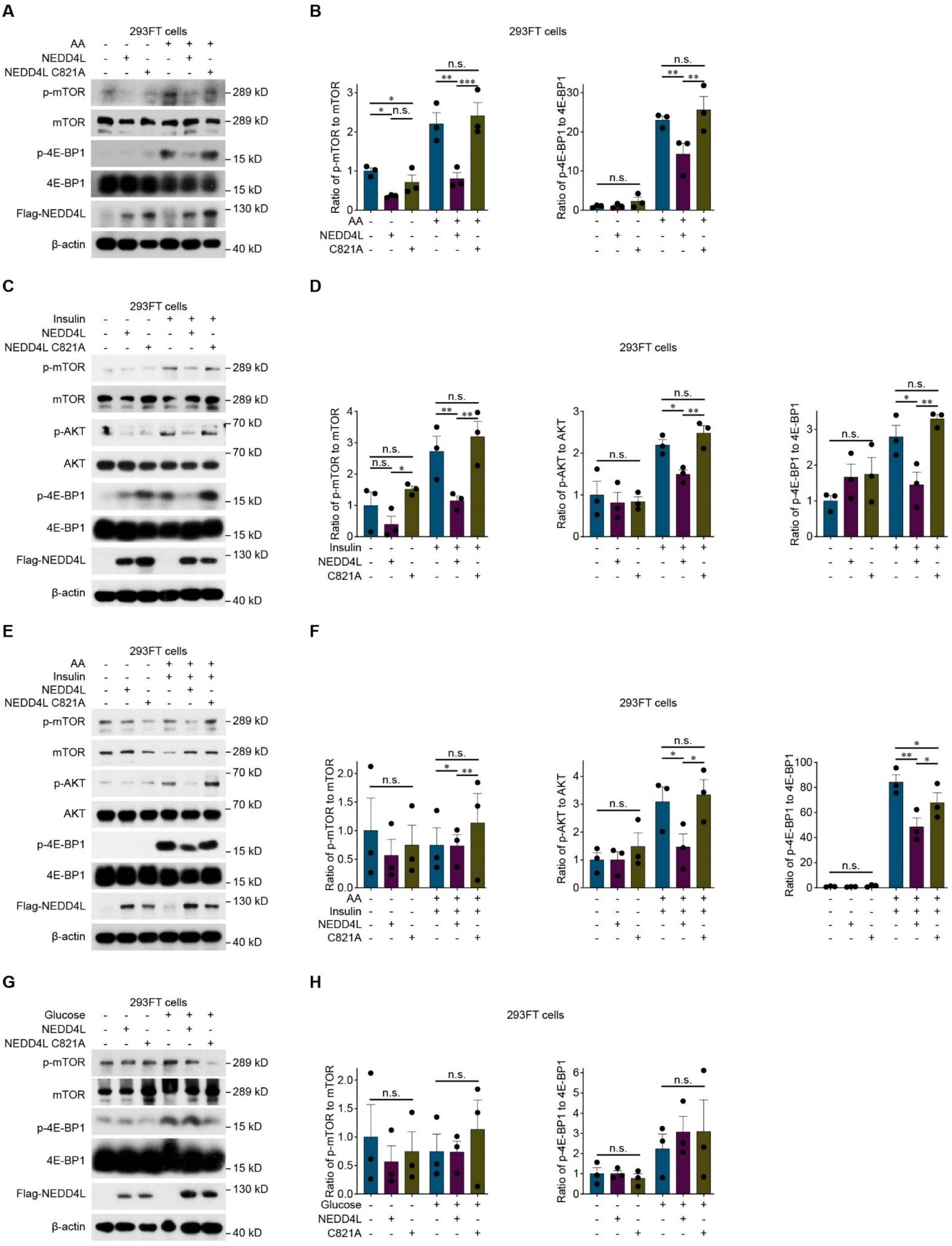
A
293FT cells
AA
NEDD4L
NEDD4L C821A
p-mTOR
mTOR
p-4E-BP1
4E-BP1
Flag-NEDD4L
β-actin
289 kD
15 kD
130 kD
40 kD
B
293FT cells
Ratio of p-mTOR to mTOR
Ratio of p-4E-BP1 to 4E-BP1
C821A
n.s.
C
Insulin
p-AKT
AKT
70 kD
D
Ratio of p-AKT to AKT
E
F
G
Glucose
H

**Figure S3**

**Figure S3.** NEDD4L inhibits the AKT/mTOR signaling pathway under stimulation with AA and insulin.

**(A, B)** Representative western blots (A) and quantification of p-mTOR levels (normalized to mTOR levels, B, left) and p-4E-BP1 levels (normalized to 4E-BP1 levels, B, right) in control 293FT cells and 293FT cells overexpressing wild-type NEDD4L (NEDD4L) or an E3 ligase activity-dead mutant of NEDD4L (C821A) incubated with or without 200 μM amino acids (AA, for 15 min). Three independent western blot analyses were performed.

**(C, D)** Representative western blots (C) and quantification of p-mTOR levels (normalized to mTOR levels, D, left), p-AKT levels (normalized to AKT levels, D, middle), and p-4E-BP1 levels (normalized to 4E-BP1 levels, D, right) in control 293FT cells and 293FT cells overexpressing wild-type NEDD4L (NEDD4L) or an E3 ligase activity-dead mutant of NEDD4L (C821A) incubated with or without 800 nM insulin (for 10 min). Three independent western blot analyses were performed.

**(E, F)** Representative western blots (E) and quantification of p-mTOR levels (normalized to mTOR levels, F, left), p-AKT levels (normalized to AKT levels, F, middle), and p-4E-BP1 levels (normalized to 4E-BP1 levels, F, right) in control 293FT cells and 293FT cells overexpressing wild-type NEDD4L (NEDD4L) or an E3 ligase activity-dead mutant of NEDD4L (C821A) incubated with or without 200 μM AA (for 15 min), and 800 nM insulin (for 10 min). Three independent western blot analyses were performed.

**(G, H)** Representative western blots (G) and quantification of p-mTOR levels (normalized to mTOR levels, H, left), and p-4E-BP1 levels (normalized to 4E-BP1 levels, H, right) in control 293FT cells and 293FT cells overexpressing wild-type NEDD4L (NEDD4L) or an E3 ligase activity-dead mutant of NEDD4L (C821A) incubated with or without 4.5 mg/ml glucose (for 20 min). Three independent western blot analyses were performed.

The data are presented as the mean ± s.e.m. values. *P*- values were determined by unpaired two-way ANOVA with uncorrected Fisher's LSD test (B, D, F, and H). * $P < 0.05$; ** $P < 0.01$; *** $P < 0.001$; n.s., not significant.

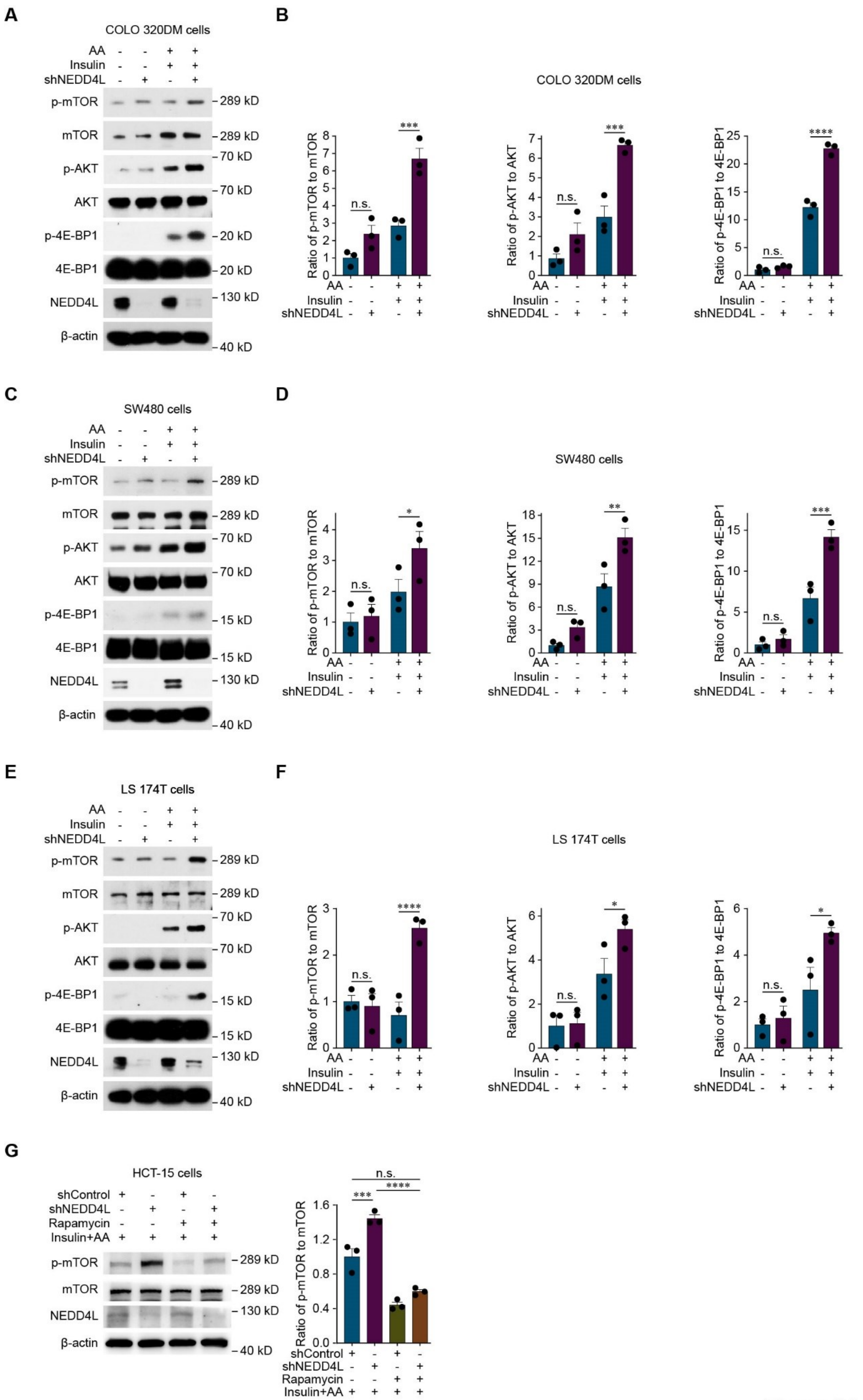
A
COLO 320DM cells
AA - - + +
Insulin - - + +
shNEDD4L - + - +
p-mTOR
mTOR
p-AKT
AKT
p-4E-BP1
4E-BP1
NEDD4L
β-actin
289 kD
289 kD
70 kD
70 kD
20 kD
20 kD
130 kD
40 kD
B
COLO 320DM cells
Ratio of p-mTOR to mTOR
Ratio of p-AKT to AKT
Ratio of p-4E-BP1 to 4E-BP1
n.s.
***
****
C
SW480 cells
p-mTOR
mTOR
p-AKT
AKT
p-4E-BP1
4E-BP1
NEDD4L
β-actin
289 kD
289 kD
70 kD
70 kD
15 kD
15 kD
130 kD
40 kD
D
SW480 cells
n.s.
*
**
***
E
LS 174T cells
p-mTOR
mTOR
p-AKT
AKT
p-4E-BP1
4E-BP1
NEDD4L
β-actin
289 kD
289 kD
70 kD
70 kD
15 kD
15 kD
130 kD
40 kD
F
LS 174T cells
n.s.
*
****
G
HCT-15 cells
shControl + - + -
shNEDD4L - + - +
Rapamycin - - + +
Insulin+AA + + + +
p-mTOR
mTOR
NEDD4L
β-actin
289 kD
289 kD
130 kD
40 kD
Ratio of p-mTOR to mTOR
n.s.
***
****

**Figure S4**

**Figure S4.** Knockdown of NEDD4L in colorectal cancer cells promotes the activation of the AKT/mTOR signaling pathway under stimulation with AA and insulin.

**(A, B)** Representative western blots (A) and quantification of p-mTOR levels (normalized to mTOR levels, B, left), p-AKT levels (normalized to AKT levels, B, middle), and p-4E-BP1 levels (normalized to 4E-BP1 levels, B, right) in control COLO 320DM cells (shControl) and NEDD4L-knockdown COLO 320DM cells (shNEDD4L) incubated with or without 200 μM amino acids (AA; for 15 min) and 800 nM insulin (for 10 min). Three independent western blot analyses were performed.

**(C, D)** Representative western blots (C) and quantification of p-mTOR levels (normalized to mTOR levels, D, left), p-AKT levels (normalized to AKT levels, D, middle), and p-4E-BP1 levels (normalized to 4E-BP1 levels, D, right) in control SW480 cells (shControl) and NEDD4L-knockdown SW480 cells (shNEDD4L) incubated with or without 200 μM AA (for 15 min) and 800 nM insulin (for 10 min). Three independent western blot analyses were performed.

**(E, F)** Representative western blots (E) and quantification of p-mTOR levels (normalized to mTOR levels, F, left), p-AKT levels (normalized to AKT levels, F, middle), and p-4E-BP1 levels (normalized to 4E-BP1 levels, F, right) in control LS 174T cells (shControl) and NEDD4L-knockdown LS 174T cells (shNEDD4L) incubated with or without 200 μM AA (for 15 min) and 800 nM insulin (for 10 min). Three independent western blot analyses were performed.

**(G)** Representative western blots (left) and quantification of p-mTOR levels (normalized to mTOR levels, right) in control HCT-15 cells (shControl) and NEDD4L-knockdown HCT-15 cells (shNEDD4L) cultured with or without 100 nM rapamycin

(24 hr) in combination with 200 μM AA (for 15 min) and 800 nM insulin (for 10 min). Three independent western blot analyses were performed.

The data are presented as the mean ± s.e.m. values. *P*- values were determined by unpaired two-way ANOVA with uncorrected Fisher's LSD test (B, D, and F), or unpaired one-way ANOVA with uncorrected Fisher's LSD test (G). * $P < 0.05$; ** $P < 0.01$; *** $P < 0.001$; **** $P < 0.0001$; n.s., not significant.

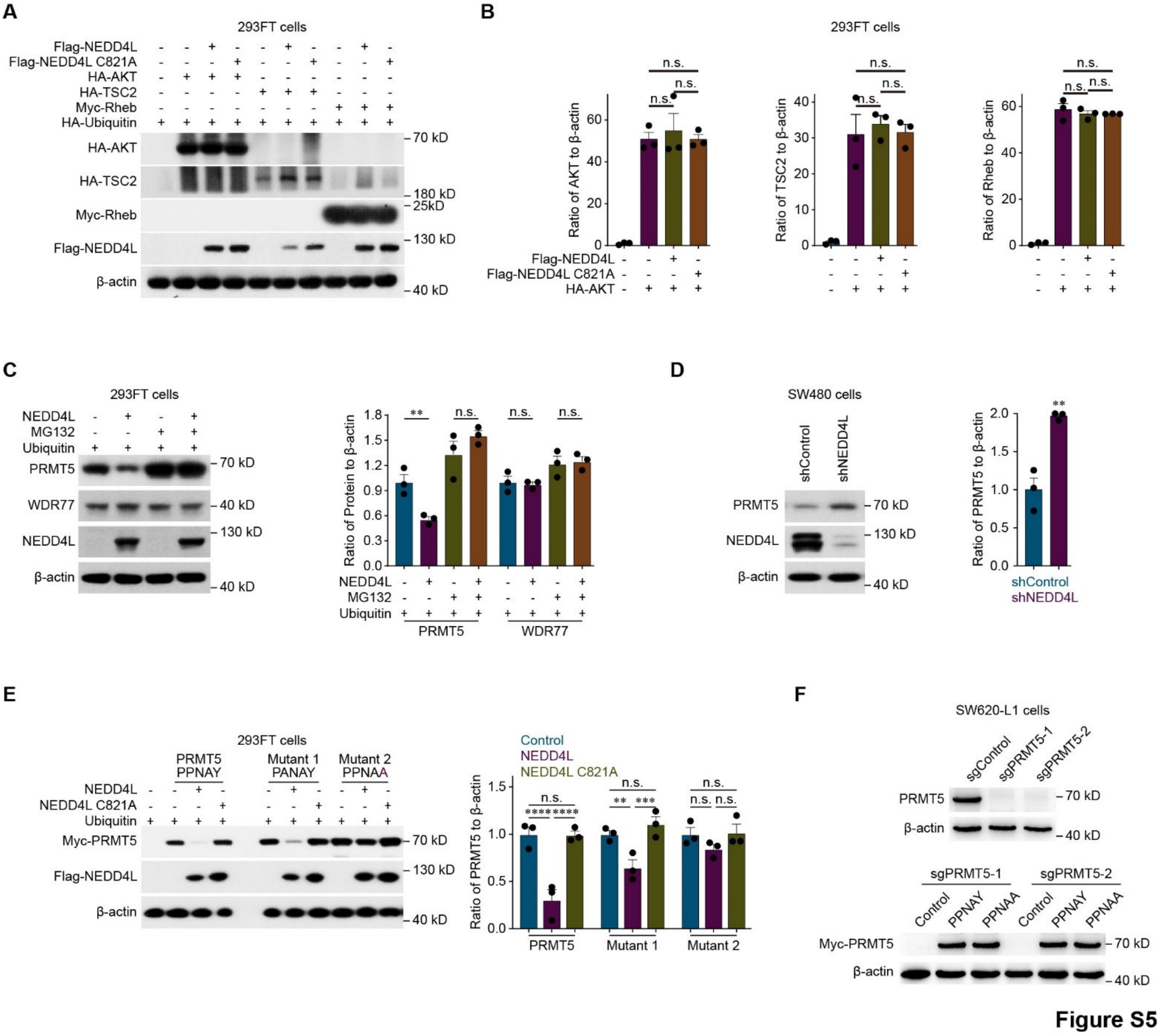


**Figure S5.** NEDD4L ubiquitinates PRMT5 to promote its degradation.

**(A, B)** Representative western blots (A) and quantification of HA-AKT, HA-TSC2, and Myc-Rheb expression (normalized to β-actin expression, B) in control 293FT cells with HA-ubiquitin overexpression and 293FT cells with overexpression of HA-ubiquitin in combination with wild-type NEDD4L (Flag-NEDD4L) or an E3 ligase activity-dead mutant of NEDD4L (NEDD4L C821A) transfected with or without HA-AKT, HA-TSC2, or Myc-Rheb. Three independent western blot analyses were performed.

**(C)** Representative western blots (left) and quantification of endogenous PRMT5 expression and WDR77 expression (normalized to β-actin expression, right) in control 293FT cells overexpressing HA-ubiquitin and 293FT cells overexpressing HA-ubiquitin in combination with wild-type NEDD4L incubated with or without 20 μM MG132 for 12 hr. Three independent western blot analyses were performed.

**(D)** Representative western blots (left) and quantification of PRMT5 expression (normalized to β-actin expression, right) in control SW480 cells (shControl) and NEDD4L-knockdown SW480 cells (shNEDD4L). Three independent western blot analyses were performed.

**(E)** Representative western blots (left) and quantification of the expression of exogenous Flag-PRMT5 containing the wild-type or mutant NEDD4L binding motif (normalized to β-actin expression, right) in 293FT cells with overexpression of wild-type NEDD4L (NEDD4L) or an E3 ligase activity-dead mutant of NEDD4L (NEDD4L C821A) in combination with ubiquitin and transfected with PRMT5 containing the wild-type NEDD4L binding motif (PRMT5, PPNAY) or one of two mutant NEDD4L binding motifs (Mutant1, PANAY; Mutant2, PPNAA). Three independent western blot analyses were performed.

**(F)** Representative western blots showing PRMT5 expression in control SW620-L1 cells (sgControl), PRMT5-knockout SW620-L1 cells (sgPRMT5-1 or sgPRMT5-2) (top), and SW620-L1 cells with endogenous PRMT5 knockout and restoration with Flag-PRMT5 containing the wild-type NEDD4L binding motif (PPNAY) or mutant NEDD4L binding motif (PPNAA) in combination with WDR77 overexpression. (bottom). Three independent western blot analyses were performed.

The data are presented as the mean ± s.e.m. values. *P*- values were determined by unpaired one-way ANOVA with uncorrected Fisher's LSD test (B), unpaired two-way ANOVA with uncorrected Fisher's LSD test (C and E), or unpaired two-tailed Student's t-test with Welch's correction (D). ** $P < 0.01$; *** $P < 0.001$; **** $P < 0.0001$; n.s., not significant.

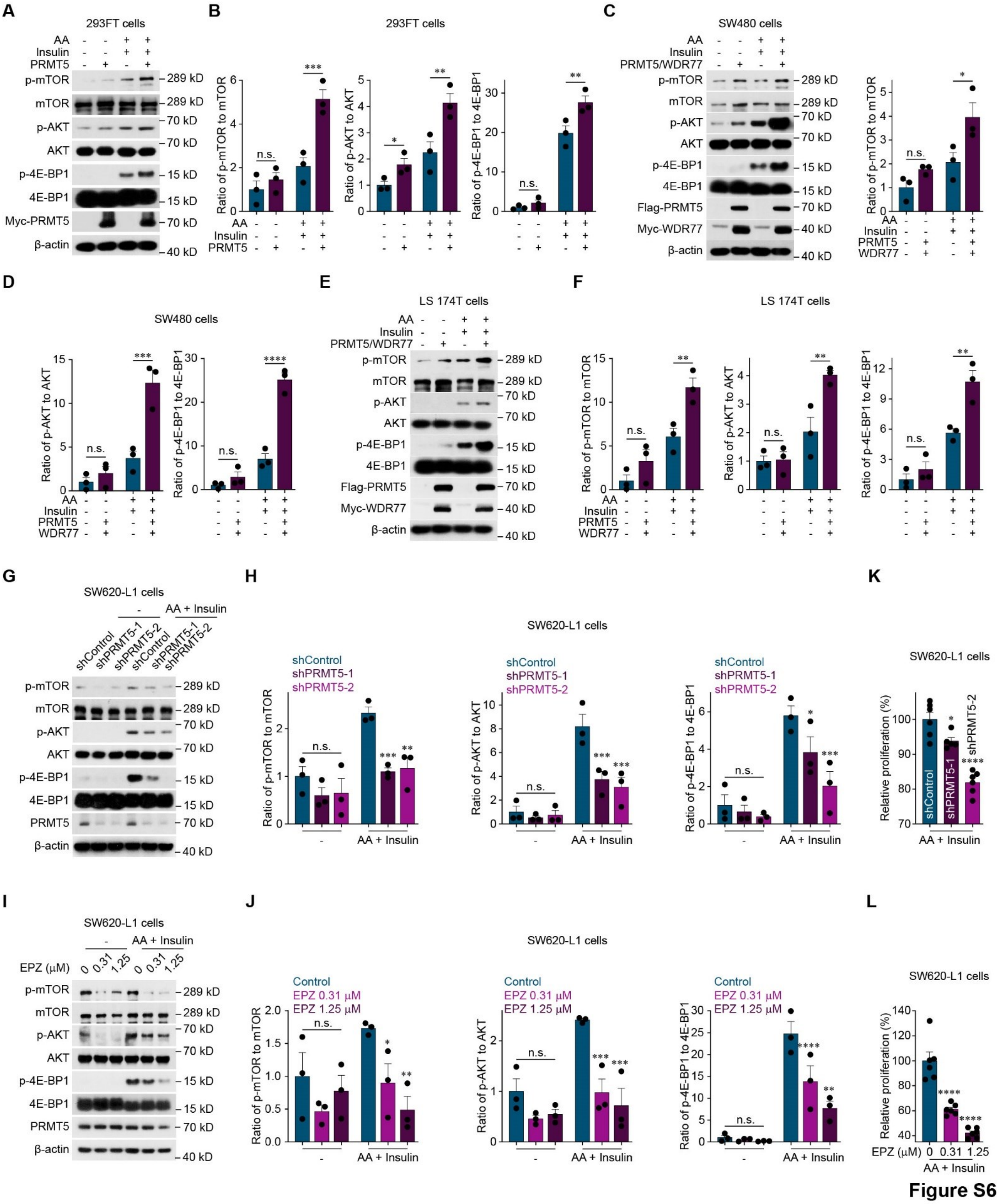


**Figure S6.** PRMT5 activates the AKT/mTOR signaling pathway to increase colorectal cancer cell proliferation.

**(A, B)** Representative western blots (A) and quantification of p-mTOR levels (normalized to mTOR levels, B, left), p-AKT levels (normalized to AKT levels, B, middle), and p-4E-BP1 levels (normalized to 4E-BP1 levels, B, right) in control 293FT cells and 293FT cells with overexpression of wild-type PRMT5 (PRMT5) incubated with or without 200 μM amino acids (AA, for 15 min) and 800 nM insulin (for 10 min). Three independent western blot analyses were performed.

**(C, D)** Representative western blots (C, left) and quantification of p-mTOR levels (normalized to mTOR levels, C, right), p-AKT levels (normalized to AKT levels, D, left), and p-4E-BP1 levels (normalized to 4E-BP1 levels, D, right) in control SW480 cells and SW480 cells with overexpression of wild-type PRMT5 in combination with WDR77 (PRMT5/WDR77) incubated with or without 200 μM AA (for 15 min) and 800 nM insulin (for 10 min). Three independent western blot analyses were performed.

**(E, F)** Representative western blots (E) and quantification of p-mTOR levels (normalized to mTOR levels, F, left), p-AKT levels (normalized to AKT levels, F, middle), and p-4E-BP1 levels (normalized by 4E-BP1 levels, F, right) in control LS 174T cells and LS 174T cells with overexpression of wild-type PRMT5 in combination with WDR77 overexpression (PRMT5/WDR77) incubated with or without 200 μM AA (for 15 min) and 800 nM insulin (for 10 min). Three independent western blot analyses were performed.

**(G, H)** Representative western blots (G) and quantification of p-mTOR levels (normalized to mTOR levels, H, left), p-AKT levels (normalized to AKT levels, H, middle), and p-4E-BP1 levels (normalized to 4E-BP1 levels, H, right) in control SW620-L1 cells (shControl), or PRMT5-knockdown SW620-L1 cells (shPRMT5-1 or shPRMT5-2) incubated with or without 200 μM AA (for 15 min) and 800 nM insulin (for 10 min). Three independent western blot analyses were performed.

**(I, J)** Representative western blots (I) and quantification of p-mTOR levels (normalized to mTOR levels, J, left), p-AKT levels (normalized to AKT levels, J, middle), and p-4E-BP1 levels (normalized to 4E-BP1 levels, J, right) in SW620-L1 cells treated with the PRMT5 inhibitor EPZ015666 (0, 0.31 or 1.25 μM, for 48 hr) and incubated with AA (for 15 min) and 800 nM insulin (for 10 min). Three independent western blot analyses were performed.

**(K)** *In vitro* proliferation assay of control SW620-L1 cells (shControl) and PRMT5-knockdown SW620-L1 cells (shPRMT5-1 or shPRMT5-2, 3,000 cells) cultured with 200 μM AA and 800 nM insulin for 24 hr. At least five independent CCK-8 assays were performed.

**(L)** *In vitro* proliferation assay of SW620-L1 cells cultured with the PRMT5 inhibitor EPZ015666 (0, 0.31 or 1.25 μM), 200 μM AA acid and 800 nM insulin for 24 hr. Six independent CCK-8 assays were performed.

The data are presented as the mean ± s.e.m. values. *P*- values were determined by unpaired two-way ANOVA with uncorrected Fisher's LSD test (B, C, D, F, H, and J), or unpaired one-way ANOVA with uncorrected Fisher's LSD test (K and L). * $P < 0.05$; ** $P < 0.01$; *** $P < 0.001$; **** $P < 0.0001$; n.s., not significant.

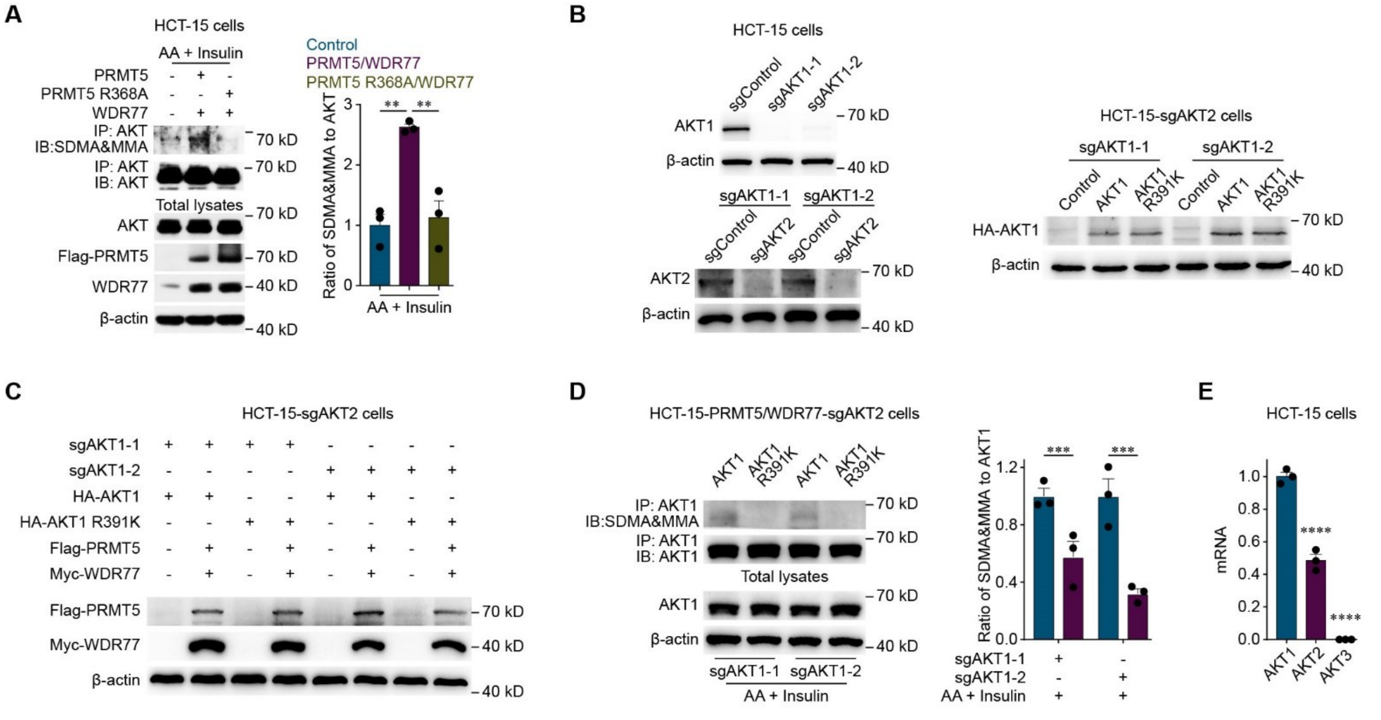


Figure S7

**Figure S7.** PRMT5 methylates R391 in AKT1.

**(A)** Representative western blots (left) and quantification of pan-AKT methylarginine levels (normalized by pan-AKT levels, right) in control HCT-15 cells, HCT-15 cells with overexpression of WDR77 in combination with overexpression of wild-type PRMT5 (PRMT5/WDR77) or a methyltransferase-inactive mutant of PRMT5 (PRMT5 R368A/WDR77) incubated with 200 μM amino acid (AA; for 15 min) and 800 nM insulin (for 10 min). Three independent western blot analyses were performed.

**(B)** Representative western blots showing AKT1 expression in control HCT-15 cells (sgControl), and AKT1 knockout HCT-15 cells (sgAKT1-1 or sgAKT1-2) (top left), AKT2 expression in control AKT1-knockout HCT-15 cells (sgControl/sgAKT1-1 or sgControl/sgAKT1-2) and AKT1 + AKT2 knockout HCT-15 cells (sgAKT2/sgAKT1-1 or sgAKT2/sgAKT1-2) (bottom left), and HA-AKT1 and HA-AKT R391K expression in control AKT1 + AKT2 knockout HCT-15 cells

(Control/sgAKT2/sgAKT1-1 or Control/sgAKT2/sgAKT1-2), HCT-15 cells with AKT1 + AKT2 knockout and restoration with wild-type AKT1 (AKT1/sgAKT2/sgAKT1-1 or AKT1/sgAKT2/sgAKT1-2) or the AKT1 R391K mutant that cannot be methylated by PRMT5 (AKT1 R391K/sgAKT2/sgAKT1-1 or AKT1 R391K /sgAKT2/sgAKT1-2) (right). Three independent western blot analyses were performed.

**(C)** Representative western blots showing Flag-PRMT5 and Myc-WDR77 expression in HCT-15 cells with Control or PRMT5 and WDR77 overexpression in combination with endogenous AKT1 + AKT2 knockout and restoration with wild-type AKT1, or the AKT1-R391K mutant that cannot be methylated by PRMT5 (right). Three independent western blot analyses were performed.

**(D)** Representative western blots (left) and quantification of AKT1 methylarginine levels (normalized to AKT1 expression, right) in HCT-15 cells with knockout of endogenous AKT2 + AKT1 combined with overexpression of PRMT5 and WDR77 (PRMT5/WDR77/sgAKT2/sgAKT1-1 or PRMT5/WDR77/sgAKT2/sgAKT1-2) and restoration with wild-type AKT1 (AKT1) or the AKT1 R391K mutant that cannot be methylated by PRMT5 (AKT R391K) incubated with 200 μM AA (for 15 min) and 800 nM insulin (for 10 min). Three independent western blot analyses were performed.

**(E)** qPCR analysis of the mRNA expression of AKT1, AKT2 and AKT3 in HCT-15 cells. Three independent experiments were performed.

The data are presented as the mean ± s.e.m. values. *P*- values were determined by unpaired one-way ANOVA with uncorrected Fisher's LSD test (B and E), or unpaired two-way ANOVA with uncorrected Fisher's LSD test (D). ** $P < 0.01$; ***, $P < 0.001$; **** $P < 0.0001$; n.s., not significant.

**Table S1.** The list of shRNAs of the shRNA library targeting 156 human E3 ubiquitin ligases.

| Gene ID | NM ID | Gene | shRNA ID |
|---|---|---|---|
| 267 | NM_001144 | AMFR | TRCN0000003373<br>TRCN0000003374<br>TRCN0000003375 |
| 10393 | NM_014885 | ANAPC10 | TRCN0000004431<br>TRCN0000004432<br>TRCN0000004433<br>TRCN0000004434<br>TRCN0000010876 |
| 51529 | NM_016476 | ANAPC11 | TRCN0000038799<br>TRCN0000038800<br>TRCN0000038801<br>TRCN0000038802<br>TRCN0000038803 |
| 29882 | NM_013366 | ANAPC2 | TRCN0000004357<br>TRCN0000004358<br>TRCN0000004359<br>TRCN0000004360<br>TRCN0000010870 |
| 29945 | NM_013367 | ANAPC4 | TRCN0000004361<br>TRCN0000004362<br>TRCN0000004363<br>TRCN0000004364<br>TRCN0000004365 |
| 51433 | NM_016237 | ANAPC5 | TRCN0000004152<br>TRCN0000004153<br>TRCN0000004154 |

| | | | |
|---|---|---|---|
| 51433 | NM_016237 | ANAPC5 | TRCN0000004155<br>TRCN0000004156 |
| 25820 | NM_005744 | ARIH1 | TRCN0000007500<br>TRCN0000007501<br>TRCN0000007502<br>TRCN0000007503<br>TRCN0000007504 |
| 580 | NM_000465 | BARD1 | TRCN0000003743<br>TRCN0000003744<br>TRCN0000003745<br>TRCN0000003746<br>TRCN0000003747 |
| 329 | NM_001166 | BIRC2 | TRCN0000003780<br>TRCN0000003781<br>TRCN0000003782<br>TRCN0000003783<br>TRCN0000003784 |
| 330 | NM_001165 | BIRC3 | TRCN0000003775<br>TRCN0000003776<br>TRCN0000003777<br>TRCN0000003778<br>TRCN0000003779 |
| 648 | NM_005180 | BMI1 | TRCN0000020154<br>TRCN0000020155<br>TRCN0000020156<br>TRCN0000020157<br>TRCN0000020158 |
| 8315 | NM_006768 | BRAP | TRCN0000007629<br>TRCN0000007630<br>TRCN0000007631 |

| | | | |
|---|---|---|---|
| 8315 | NM_006768 | BRAP | TRCN0000007632<br>TRCN0000007633 |
| 672 | NM_007294 | BRCA1 | TRCN0000009823<br>TRCN0000009824<br>TRCN0000010305<br>TRCN0000039833<br>TRCN0000039834<br>TRCN0000039835<br>TRCN0000039836<br>TRCN0000039837 |
| 8945 | NM_003939 | BTRC | TRCN0000006541<br>TRCN0000006542<br>TRCN0000006543<br>TRCN0000006544<br>TRCN0000006545 |
| 283450 | NM_173813 | C12orf51 | TRCN0000004992<br>TRCN0000004993<br>TRCN0000004994<br>TRCN0000004995<br>TRCN0000004996 |
| 79596 | NM_024546 | C13orf7 | TRCN0000133667<br>TRCN0000134953<br>TRCN0000136897<br>TRCN0000137082<br>TRCN0000137111 |
| 55148 | NM_018108 | C14orf130 | TRCN0000037024<br>TRCN0000037025<br>TRCN0000037026<br>TRCN0000037027<br>TRCN0000037028 |

| | | | |
|---|---|---|---|
| 79594 | NM_024544 | C1orf166 | TRCN0000033934<br>TRCN0000033936<br>TRCN0000033937<br>TRCN0000033938 |
| 55832 | NM_018448 | CAND1 | TRCN0000003459<br>TRCN0000003460<br>TRCN0000003461<br>TRCN0000003462<br>TRCN0000003463 |
| 867 | NM_005188 | CBL | TRCN0000010310<br>TRCN0000010311<br>TRCN0000039723<br>TRCN0000039724<br>TRCN0000039725<br>TRCN0000039726<br>TRCN0000039727 |
| 57332 | NM_020649 | CBX8 | TRCN0000021894<br>TRCN0000021895<br>TRCN0000021896<br>TRCN0000021897<br>TRCN0000021898 |
| 8697 | NM_004661 | CDC23 | TRCN0000007375<br>TRCN0000007376<br>TRCN0000007377<br>TRCN0000007378<br>TRCN0000007379 |
| 55743 | NM_018223 | CHFR | TRCN0000007702<br>TRCN0000007703<br>TRCN0000007704<br>TRCN0000007705 |

| | | | |
|---|---|---|---|
| 55743 | NM_018223 | CHFR | TRCN0000007706 |
| 4850 | NM_013316 | CNOT4 | TRCN0000015213<br>TRCN0000015214<br>TRCN0000015215<br>TRCN0000015216<br>TRCN0000015217 |
| 8454 | NM_003592 | CUL1 | TRCN0000003391<br>TRCN0000003392<br>TRCN0000003393<br>TRCN0000003394<br>TRCN0000010781 |
| 8453 | NM_003591 | CUL2 | TRCN0000006522<br>TRCN0000006523<br>TRCN0000006524<br>TRCN0000006525<br>TRCN0000006526 |
| 8452 | NM_003590 | CUL3 | TRCN0000073343<br>TRCN0000073344<br>TRCN0000073345<br>TRCN0000073346<br>TRCN0000073347 |
| 8451 | NM_003589 | CUL4A | TRCN0000006527<br>TRCN0000006528<br>TRCN0000006529<br>TRCN0000006530<br>TRCN0000006531 |
| 8450 | NM_003588 | CUL4B | TRCN0000006532<br>TRCN0000006533<br>TRCN0000006534<br>TRCN0000006535 |

| | | | |
|---|---|---|---|
| 8450 | NM_003588 | CUL4B | TRCN0000006536 |
| 8065 | NM_003478 | CUL5 | TRCN0000006537<br>TRCN0000006538<br>TRCN0000006539<br>TRCN0000006540<br>TRCN0000011030 |
| 9820 | NM_014780 | CUL7 | TRCN0000006480<br>TRCN0000006481<br>TRCN0000006482<br>TRCN0000006483<br>TRCN0000006484 |
| 1643 | NM_000107 | DDB2 | TRCN0000083993<br>TRCN0000083994<br>TRCN0000083995<br>TRCN0000083996<br>TRCN0000083997 |
| 113878 | NM_020892 | DTX2 | TRCN0000004558<br>TRCN0000004559<br>TRCN0000004560<br>TRCN0000004561 |
| 9666 | NM_014648 | DZIP3 | TRCN0000034244<br>TRCN0000034246<br>TRCN0000034247<br>TRCN0000034248 |
| 51366 | NM_015902 | EDD1 | TRCN0000003408<br>TRCN0000003409<br>TRCN0000003410<br>TRCN0000003411<br>TRCN0000003412 |
| 2033 | NM_001429 | EP300 | TRCN0000009882 |

| | | | |
|---|---|---|---|
| 2033 | NM_001429 | EP300 | TRCN0000009883<br>TRCN0000009884<br>TRCN0000039883<br>TRCN0000039884<br>TRCN0000039885<br>TRCN0000039886<br>TRCN0000039887 |
| 1161 | NM_000082 | ERCC8 | TRCN0000003719<br>TRCN0000003720<br>TRCN0000003721<br>TRCN0000003722<br>TRCN0000003723 |
| 55120 | NM_018062 | FANCL | TRCN0000083298<br>TRCN0000083299<br>TRCN0000083300<br>TRCN0000083301<br>TRCN0000083302 |
| 25827 | NM_012157 | FBXL2 | TRCN0000004276<br>TRCN0000004277<br>TRCN0000004278<br>TRCN0000004279<br>TRCN0000004280 |
| 26223 | NM_012159 | FBXL21 | TRCN0000004286<br>TRCN0000004287<br>TRCN0000004289<br>TRCN0000010866 |
| 26224 | NM_012158 | FBXL3 | TRCN0000004281<br>TRCN0000004282<br>TRCN0000004283<br>TRCN0000004284 |

| | | | |
|---|---|---|---|
| 26224 | NM_012158 | FBXL3 | TRCN0000004285 |
| | | | TRCN0000004290 |
| | | | TRCN0000004291 |
| 26234 | NM_012161 | FBXL5 | TRCN0000004292 |
| | | | TRCN0000004293 |
| | | | TRCN0000004294 |
| | | | TRCN0000004295 |
| | | | TRCN0000004296 |
| 26233 | NM_012162 | FBXL6 | TRCN0000004297 |
| | | | TRCN0000004298 |
| | | | TRCN0000004299 |
| | | | TRCN0000118322 |
| | | | TRCN0000118323 |
| 23194 | NM_012304 | FBXL7 | TRCN0000118324 |
| | | | TRCN0000118325 |
| | | | TRCN0000118326 |
| | | | TRCN0000004300 |
| | | | TRCN0000004301 |
| 80204 | NM_012167 | FBXO11 | TRCN0000004302 |
| | | | TRCN0000004303 |
| | | | TRCN0000004304 |
| | | | TRCN0000004305 |
| | | | TRCN0000004306 |
| 26232 | NM_012168 | FBXO2 | TRCN0000004307 |
| | | | TRCN0000010867 |
| | | | TRCN0000010868 |
| | | | TRCN0000034284 |
| | | | TRCN0000034285 |
| 23014 | NM_015002 | FBXO21 | TRCN0000034286 |
| | | | TRCN0000034287 |

| | | | |
|---|---|---|---|
| 23014 | NM_015002 | FBXO21 | TRCN0000034288 |
| | | | TRCN0000004308 |
| | | | TRCN0000004309 |
| 26263 | NM_012170 | FBXO22 | TRCN0000004310 |
| | | | TRCN0000004311 |
| | | | TRCN0000010869 |
| | | | TRCN0000004312 |
| | | | TRCN0000004313 |
| 26261 | NM_012172 | FBXO24 | TRCN0000004314 |
| | | | TRCN0000004315 |
| | | | TRCN0000004316 |
| | | | TRCN0000004317 |
| | | | TRCN0000004318 |
| 26260 | NM_012173 | FBXO25 | TRCN0000004319 |
| | | | TRCN0000004320 |
| | | | TRCN0000004321 |
| | | | TRCN0000004327 |
| | | | TRCN0000004328 |
| 26273 | NM_012175 | FBXO3 | TRCN0000004329 |
| | | | TRCN0000004330 |
| | | | TRCN0000004331 |
| | | | TRCN0000034319 |
| | | | TRCN0000034320 |
| 26272 | NM_012176 | FBXO4 | TRCN0000034321 |
| | | | TRCN0000034322 |
| | | | TRCN0000034323 |
| | | | TRCN0000007731 |
| | | | TRCN0000007732 |
| 26270 | NM_018438 | FBXO6 | TRCN0000007733 |
| | | | TRCN0000007734 |

| | | | |
|---|---|---|---|
| 26270 | NM_018438 | FBXO6 | TRCN0000011101 |
| 25793 | NM_012179 | FBXO7 | TRCN0000004337<br>TRCN0000004338<br>TRCN0000004339<br>TRCN0000004340<br>TRCN0000004341 |
| 26268 | NM_012347 | FBXO9 | TRCN0000034309<br>TRCN0000034310<br>TRCN0000034311<br>TRCN0000034312<br>TRCN0000034313 |
| 23291 | NM_012300 | FBXW11 | TRCN0000004342<br>TRCN0000004343<br>TRCN0000004344<br>TRCN0000004345<br>TRCN0000004346 |
| 26190 | NM_012164 | FBXW2 | TRCN0000006546<br>TRCN0000006547<br>TRCN0000006548<br>TRCN0000006549<br>TRCN0000006550 |
| 55527 | NM_018708 | FEM1A | TRCN0000060758<br>TRCN0000060759<br>TRCN0000060760<br>TRCN0000060761 |
| 10116 | NM_015322 | FEM1B | TRCN0000060579<br>TRCN0000060580<br>TRCN0000060581<br>TRCN0000060582 |
| 57531 | NM_020771 | HACE1 | TRCN0000003413 |

| | | | |
|---|---|---|---|
| 57531 | NM_020771 | HACE1 | TRCN0000003414<br>TRCN0000003415<br>TRCN0000003416<br>TRCN0000003417 |
| 25831 | NM_015382 | HECTD1 | TRCN0000004083<br>TRCN0000004084<br>TRCN0000004085<br>TRCN0000004086<br>TRCN0000004087 |
| 143279 | NM_173497 | HECTD2 | TRCN0000007758<br>TRCN0000007759<br>TRCN0000007760<br>TRCN0000007761<br>TRCN0000007762 |
| 79654 | XM_371246 | HECTD3 | TRCN0000118242<br>TRCN0000118243<br>TRCN0000118244<br>TRCN0000118245<br>TRCN0000118246 |
| 23072 | NM_015052 | HECW1 | TRCN0000001523<br>TRCN0000001524<br>TRCN0000010637<br>TRCN0000010638<br>TRCN0000010639 |
| 57520 | XM_038999 | HECW2 | TRCN0000004789<br>TRCN0000004790<br>TRCN0000004791<br>TRCN0000004792<br>TRCN0000004793 |
| 8925 | NM_003922 | HERC1 | TRCN0000007243 |

| | | | |
|---|---|---|---|
| 8925 | NM_003922 | HERC1 | TRCN0000007244<br>TRCN0000007245<br>TRCN0000007246<br>TRCN0000007247 |
| 8924 | NM_004667 | HERC2 | TRCN0000007380<br>TRCN0000007381<br>TRCN0000007382<br>TRCN0000007383<br>TRCN0000007384 |
| 8916 | NM_014606 | HERC3 | TRCN0000000291<br>TRCN0000000292<br>TRCN0000000293<br>TRCN0000000294<br>TRCN0000000295 |
| 26091 | NM_015601 | HERC4 | TRCN0000034299<br>TRCN0000034300<br>TRCN0000034301<br>TRCN0000034302<br>TRCN0000034303 |
| 51191 | NM_016323 | HERC5 | TRCN0000004168<br>TRCN0000004169<br>TRCN0000004170<br>TRCN0000004171<br>TRCN0000010859 |
| 55008 | NM_017912 | HERC6 | TRCN0000158995<br>TRCN0000160017<br>TRCN0000160044<br>TRCN0000160299 |
| 10075 | NM_031407 | HUWE1 | TRCN0000073303<br>TRCN0000073304 |

| | | | |
|---|---|---|---|
| 10075 | NM_031407 | HUWE1 | TRCN0000073305<br>TRCN0000073306<br>TRCN0000073307 |
| 154214 | NM_152553 | IBRDC1 | TRCN0000034124<br>TRCN0000034125<br>TRCN0000034126<br>TRCN0000034127<br>TRCN0000034128 |
| 255488 | NM_182757 | IBRDC2 | TRCN0000034149<br>TRCN0000034150<br>TRCN0000034151<br>TRCN0000034152<br>TRCN0000034153 |
| 83737 | NM_031483 | ITCH | TRCN0000002087<br>TRCN0000002088<br>TRCN0000002089<br>TRCN0000002090<br>TRCN0000010680 |
| 9870 | NM_014821 | KIAA0317 | TRCN0000004423<br>TRCN0000004424<br>TRCN0000004425<br>TRCN0000004426<br>TRCN0000010874 |
| 55632 | NM_017769 | KIAA1333 | TRCN0000004220<br>TRCN0000004221<br>TRCN0000004222<br>TRCN0000004223<br>TRCN0000004224 |
| 4008 | NM_005358 | LMO7 | TRCN0000006489<br>TRCN0000006490 |

| | | | |
|---|---|---|---|
| 4008 | NM_005358 | LMO7 | TRCN0000006491<br>TRCN0000006492<br>TRCN0000006493 |
| 222484 | NM_153371 | LNX2 | TRCN0000007745<br>TRCN0000007746<br>TRCN0000007747<br>TRCN0000007748<br>TRCN0000011197 |
| 164832 | NM_198461 | LONRF2 | TRCN0000007825<br>TRCN0000007826<br>TRCN0000007827<br>TRCN0000007828<br>TRCN0000007829 |
| 79836 | NM_024778 | LONRF3 | TRCN0000022424<br>TRCN0000022425<br>TRCN0000022426<br>TRCN0000022427<br>TRCN0000022428 |
| 10892 | NM_006785 | MALT1 | TRCN0000073823<br>TRCN0000073824<br>TRCN0000073825<br>TRCN0000073826<br>TRCN0000222552 |
| 64844 | NM_022826 | MARCH7 | TRCN0000073068<br>TRCN0000073070<br>TRCN0000073072<br>TRCN0000222575<br>TRCN0000222576 |
| 57574 | NM_020814 | MARCHF4 | TRCN0000127530<br>TRCN0000129384 |

| | | | |
|---|---|---|---|
| 57574 | NM_020814 | MARCHF4 | TRCN0000130947<br>TRCN0000130948<br>TRCN0000130962 |
| 54708 | NM_017824 | MARCHF5 | TRCN0000037014<br>TRCN0000037015<br>TRCN0000037016<br>TRCN0000037017<br>TRCN0000037018 |
| 92979 | NM_138396 | MARCHF9 | TRCN0000073168<br>TRCN0000073169<br>TRCN0000073170<br>TRCN0000073171<br>TRCN0000073172 |
| 4193 | NM_002392 | MDM2 | TRCN0000003376<br>TRCN0000003377<br>TRCN0000003378<br>TRCN0000003379<br>TRCN0000003380 |
| 112950 | NM_001001651 | MED8 | TRCN0000052948<br>TRCN0000052949<br>TRCN0000052950<br>TRCN0000052951<br>TRCN0000052952<br>TRCN0000174212 |
| 23295 | NM_015246 | MGRN1 | TRCN0000033789<br>TRCN0000033790<br>TRCN0000033791<br>TRCN0000033792<br>TRCN0000033793 |
| 54542 | NM_018835 | MNAB | TRCN0000037009 |

| | | | |
|---|---|---|---|
| 54542 | NM_018835 | MNAB | TRCN0000037012<br>TRCN0000037013 |
| 29116 | NM_013262 | MYLIP | TRCN0000033819<br>TRCN0000033820<br>TRCN0000033821<br>TRCN0000033822<br>TRCN0000033823 |
| 4734 | NM_006154 | NEDD4 | TRCN0000007550<br>TRCN0000007551<br>TRCN0000007552<br>TRCN0000007553<br>TRCN0000007554 |
| 23327 | NM_015277 | NEDD4L | TRCN0000000904<br>TRCN0000000905<br>TRCN0000000906<br>TRCN0000000907<br>TRCN0000000908 |
| 51070 | NM_015953 | NOSIP | TRCN0000045603<br>TRCN0000045604<br>TRCN0000045605<br>TRCN0000045606<br>TRCN0000045607 |
| 5071 | NM_013988 | PARK2 | TRCN0000000281<br>TRCN0000000282<br>TRCN0000000283<br>TRCN0000000284<br>TRCN0000000285 |
| 84108 | NM_032154 | PCGF6 | TRCN0000073108<br>TRCN0000073109<br>TRCN0000073110 |

| | | | |
|---|---|---|---|
| 84108 | NM_032154 | PCGF6 | TRCN0000073111<br>TRCN0000073112 |
| 23759 | NM_014337 | PPIL2 | TRCN0000000160<br>TRCN0000000161<br>TRCN0000000162<br>TRCN0000000163<br>TRCN0000000164 |
| 27339 | NM_014502 | PRPF19 | TRCN0000006592<br>TRCN0000006593<br>TRCN0000006594<br>TRCN0000006595<br>TRCN0000006596 |
| 5930 | NM_006910 | RBBP6 | TRCN0000034214<br>TRCN0000034215<br>TRCN0000034216<br>TRCN0000034217<br>TRCN0000034218 |
| 149041 | NM_172071 | RC3H1 | TRCN0000122428<br>TRCN0000122593<br>TRCN0000122891<br>TRCN0000139015<br>TRCN0000139513<br>TRCN0000139559<br>TRCN0000140092<br>TRCN0000142153 |
| 149041 | NM_172071 | RC3H1 | TRCN0000142634<br>TRCN0000144045 |
| 6015 | NM_002931 | RING1 | TRCN0000021989<br>TRCN0000021990<br>TRCN0000021991 |

| | | | |
|---|---|---|---|
| 6015 | NM_002931 | RING1 | TRCN0000021992<br>TRCN0000021993 |
| 26994 | NM_014372 | RNF11 | TRCN0000038794<br>TRCN0000038795<br>TRCN0000038796<br>TRCN0000038797<br>TRCN0000038798 |
| 51132 | NM_016120 | RNF12 | TRCN0000004139<br>TRCN0000004140<br>TRCN0000004141<br>TRCN0000004142<br>TRCN0000004143 |
| 55298 | NM_018320 | RNF121 | TRCN0000007721<br>TRCN0000007722<br>TRCN0000007723<br>TRCN0000007724<br>TRCN0000007725 |
| 54941 | NM_017831 | RNF125 | TRCN0000004230<br>TRCN0000004231<br>TRCN0000004232<br>TRCN0000004233<br>TRCN0000004234 |
| 79589 | NM_024539 | RNF128 | TRCN0000004794<br>TRCN0000004795<br>TRCN0000004796<br>TRCN0000004797<br>TRCN0000004798 |
| 55819 | NM_018434 | RNF130 | TRCN0000007726<br>TRCN0000007727<br>TRCN0000007728 |

| | | | |
|---|---|---|---|
| 55819 | NM_018434 | RNF130 | TRCN0000007729<br>TRCN0000007730 |
| 168433 | NM_139175 | RNF133 | TRCN0000011147<br>TRCN0000011148<br>TRCN0000011149<br>TRCN0000011150<br>TRCN0000011151 |
| 9604 | NM_004290 | RNF14 | TRCN0000003442<br>TRCN0000003443<br>TRCN0000003444<br>TRCN0000010785<br>TRCN0000010786 |
| 378925 | NM_198085 | RNF148 | TRCN0000004799<br>TRCN0000004800<br>TRCN0000004801<br>TRCN0000004802<br>TRCN0000004803 |
| 284996 | NM_173647 | RNF149 | TRCN0000034154<br>TRCN0000034155<br>TRCN0000034156<br>TRCN0000034157<br>TRCN0000034158 |
| 57484 | NM_020724 | RNF150 | TRCN0000118552<br>TRCN0000118553<br>TRCN0000118554<br>TRCN0000118555<br>TRCN0000118556 |
| 220441 | NM_173557 | RNF152 | TRCN0000007763<br>TRCN0000007764<br>TRCN0000007765 |

| | | | |
|---|---|---|---|
| 220441 | NM_173557 | RNF152 | TRCN0000007766<br>TRCN0000007767 |
| 26001 | NM_015528 | RNF167 | TRCN0000004097<br>TRCN0000004098<br>TRCN0000004099<br>TRCN0000004100<br>TRCN0000004101 |
| 285533 | NM_173662 | RNF175 | TRCN0000007768<br>TRCN0000007769<br>TRCN0000007770<br>TRCN0000007771<br>TRCN0000007772 |
| 54546 | NM_019062 | RNF186 | TRCN0000004502<br>TRCN0000004503<br>TRCN0000004504<br>TRCN0000004505<br>TRCN0000004506 |
| 25897 | NM_015435 | RNF19 | TRCN0000004804<br>TRCN0000004805<br>TRCN0000004806<br>TRCN0000004807<br>TRCN0000004808 |
| 162333 | NM_152598 | RNF190 | TRCN0000073213<br>TRCN0000073214<br>TRCN0000073215<br>TRCN0000073216<br>TRCN0000073217 |
| 6045 | NM_007212 | RNF2 | TRCN0000033694<br>TRCN0000033695<br>TRCN0000033696 |

| | | | |
|---|---|---|---|
| 6045 | NM_007212 | RNF2 | TRCN0000033697<br>TRCN0000033698 |
| 56254 | NM_019592 | RNF20 | TRCN0000033874<br>TRCN0000033875<br>TRCN0000033876<br>TRCN0000033877<br>TRCN0000033878 |
| 9810 | NM_014771 | RNF40 | TRCN0000004780<br>TRCN0000004781<br>TRCN0000004782<br>TRCN0000004783<br>TRCN0000004784 |
| 9616 | NM_014245 | RNF7 | TRCN0000038804<br>TRCN0000038805<br>TRCN0000038806<br>TRCN0000038807<br>TRCN0000038808 |
| 9025 | NM_003958 | RNF8 | TRCN0000003437<br>TRCN0000003438<br>TRCN0000003439<br>TRCN0000003440<br>TRCN0000003441 |
| 6502 | NM_005983 | SKP2 | TRCN0000007530<br>TRCN0000007531<br>TRCN0000007532<br>TRCN0000007533<br>TRCN0000007534 |
| 57154 | NM_020429 | SMURF1 | TRCN0000003471<br>TRCN0000003472<br>TRCN0000003473 |

| | | | |
|---|---|---|---|
| 57154 | NM_020429 | SMURF1 | TRCN0000003474<br>TRCN0000010791 |
| 64750 | NM_022739 | SMURF2 | TRCN0000003475<br>TRCN0000003476<br>TRCN0000003477<br>TRCN0000003478<br>TRCN0000010792 |
| 10273 | NM_005861 | STUB1 | TRCN0000007525<br>TRCN0000007526<br>TRCN0000007527<br>TRCN0000007528<br>TRCN0000007529 |
| 6921 | NM_005648 | TCEB1 | TRCN0000022124<br>TRCN0000022125<br>TRCN0000022126<br>TRCN0000022127<br>TRCN0000022128 |
| 6923 | NM_007108 | TCEB2 | TRCN0000007661<br>TRCN0000007662<br>TRCN0000011096<br>TRCN0000011097<br>TRCN0000011098 |
| 7188 | NM_004619 | TRAF5 | TRCN0000007343<br>TRCN0000007344<br>TRCN0000007345<br>TRCN0000007346<br>TRCN0000007347 |
| 7189 | NM_004620 | TRAF6 | TRCN0000007348<br>TRCN0000007349<br>TRCN0000007350 |

| | | | |
|---|---|---|---|
| 7189 | NM_004620 | TRAF6 | TRCN0000007351<br>TRCN0000007352 |
| 84231 | NM_032271 | TRAF7 | TRCN0000056948<br>TRCN0000056949<br>TRCN0000056950<br>TRCN0000056951<br>TRCN0000056952 |
| 54476 | NM_019011 | TRIAD3 | TRCN0000003464<br>TRCN0000003465<br>TRCN0000003466<br>TRCN0000010788<br>TRCN0000010789 |
| 373 | NM_001656 | TRIM23 | TRCN0000034204<br>TRCN0000034205<br>TRCN0000034206<br>TRCN0000034208 |
| 22954 | NM_012210 | TRIM32 | TRCN0000003455<br>TRCN0000003456<br>TRCN0000003457<br>TRCN0000003458<br>TRCN0000010787 |
| 79097 | NM_024114 | TRIM48 | TRCN0000033919<br>TRCN0000033920<br>TRCN0000033921<br>TRCN0000033922<br>TRCN0000033923 |
| 57093 | NM_020358 | TRIM49 | TRCN0000033884<br>TRCN0000033885<br>TRCN0000033886<br>TRCN0000033887 |

| | | | |
|---|---|---|---|
| 57093 | NM_020358 | TRIM49 | TRCN0000033888 |
| 84676 | NM_032588 | TRIM63 | TRCN0000073118 |
| | | | TRCN0000073119 |
| | | | TRCN0000073120 |
| | | | TRCN0000073121 |
| | | | TRCN0000073122 |
| 9320 | NM_004238 | TRIP12 | TRCN0000022374 |
| | | | TRCN0000022375 |
| | | | TRCN0000022376 |
| | | | TRCN0000022377 |
| | | | TRCN0000022378 |
| 26262 | NM_130465 | TSPAN17 | TRCN0000034304 |
| | | | TRCN0000034305 |
| | | | TRCN0000034306 |
| | | | TRCN0000034307 |
| | | | TRCN0000034308 |
| 7337 | NM_000462 | UBE3A | TRCN0000003368 |
| | | | TRCN0000003369 |
| | | | TRCN0000003370 |
| | | | TRCN0000003371 |
| | | | TRCN0000003372 |
| 89910 | NM_130466 | UBE3B | TRCN0000004775 |
| | | | TRCN0000004776 |
| | | | TRCN0000004777 |
| | | | TRCN0000004778 |
| | | | TRCN0000004779 |
| 9690 | NM_014671 | UBE3C | TRCN0000003399 |
| | | | TRCN0000003400 |
| | | | TRCN0000003401 |
| | | | TRCN0000003402 |

| | | | |
|---|---|---|---|
| 9690 | NM_014671 | UBE3C | TRCN0000010783 |
| 9354 | NM_004788 | UBE4A | TRCN0000007395<br>TRCN0000007396<br>TRCN0000007397<br>TRCN0000007398<br>TRCN0000007399 |
| 10277 | NM_006048 | UBE4B | TRCN0000007545<br>TRCN0000007546<br>TRCN0000007547<br>TRCN0000007548<br>TRCN0000007549 |
| 22888 | NM_014948 | UBOX5 | TRCN0000004437<br>TRCN0000004438<br>TRCN0000004439<br>TRCN0000004440<br>TRCN0000004441 |
| 197131 | NM_174916 | UBR1 | TRCN0000003423<br>TRCN0000003424<br>TRCN0000003425<br>TRCN0000003426<br>TRCN0000003427 |
| 23304 | NM_015255 | UBR2 | TRCN0000003403<br>TRCN0000003404<br>TRCN0000003405<br>TRCN0000003406<br>TRCN0000003407 |
| 23352 | NM_020765 | UBR4 | TRCN0000154749<br>TRCN0000155927<br>TRCN0000157658<br>TRCN0000157737 |

| | | | |
|---|---|---|---|
| 23352 | NM_020765 | UBR4 | TRCN0000152115<br>TRCN0000154429<br>TRCN0000154886<br>TRCN0000155202<br>TRCN0000155617 |
| 115426 | NM_152306 | UHRF2 | TRCN0000003479<br>TRCN0000003480<br>TRCN0000003481<br>TRCN0000003482<br>TRCN0000010793 |
| 7428 | NM_000551 | VHL | TRCN0000010459<br>TRCN0000010460<br>TRCN0000010461<br>TRCN0000039623<br>TRCN0000039624<br>TRCN0000039625<br>TRCN0000039626<br>TRCN0000039627 |
| 151525 | NM_152528 | WDSUB1 | TRCN0000073203<br>TRCN0000073204<br>TRCN0000073205<br>TRCN0000073206<br>TRCN0000073207 |
| 11059 | NM_007013 | WWP1 | TRCN0000003395<br>TRCN0000003396<br>TRCN0000003397<br>TRCN0000003398<br>TRCN0000010782 |
| 11060 | NM_007014 | WWP2 | TRCN0000001512<br>TRCN0000001513 |

| | | | |
|---|---|---|---|
| 11060 | NM_007014 | WWP2 | TRCN0000001514<br>TRCN0000001515<br>TRCN0000001516 |
| 10444 | NM_006336 | ZER1 | TRCN0000139495<br>TRCN0000139522<br>TRCN0000140548<br>TRCN0000140793<br>TRCN0000143404<br>TRCN0000121635<br>TRCN0000122249 |
| 10444 | NM_006336 | ZER1 | TRCN0000144077<br>TRCN0000144439<br>TRCN0000144625 |
| 130507 | NM_172070 | ZNF650 | TRCN0000034099<br>TRCN0000034100<br>TRCN0000034101<br>TRCN0000034102<br>TRCN0000034103 |

**Table S2.** Mass spectrometry results for the series coimmunoprecipitated proteins associated with the Flag-NEDD4L and HA-Ubiquitin.

| Uniprot accession | Protein name | kDa | Score | Unique peptides |
|---|---|---|---|---|
| Q4VCS5 | Angiomotin (AMOT) | 118.47 | 936 | 30 |
| Q9BQA1 | Methylosome protein 50 (WDR77) | 37.44 | 344 | 8 |
| **O14744** | **Protein arginine N-methyltransferase 5 (PRMT5)** | **73.32** | **335** | **14** |
| P46934 | E3 ubiquitin-protein ligase NEDD4 (NEDD4) | 150.28 | 322 | 7 |
| P0CG47 | Polyubiquitin-B (UBB) | 25.80 | 229 | 6 |
| Q8IY63 | Angiomotin-like protein 1 (AMOTL1) | 106.85 | 219 | 11 |
| P15924 | Desmoplakin (DSP) | 334.02 | 163 | 6 |
| P62258 | 14-3-3 protein epsilon (YWHAE) | 29.32 | 94 | 3 |
| Q969T9 | WW domain-binding protein 2 (WBP2) | 28.18 | 76 | 3 |
| Q15018 | BRISC complex subunit Abro1 (FAM175B) | 47.1 | 62 | 2 |
| Q9HCE7 | E3 ubiquitin-protein ligase SMURF1 (SMURF1) | 86.92 | 57 | 2 |
| Q9NV92 | NEDD4 family-interacting protein 2 (NDFIP2) | 36.71 | 52 | 2 |
| P07355 | Annexin A2 (ANXA2) | 38.81 | 43 | 3 |
| Q8TER0 | Sushi, nidogen and EGF-like domain-containing protein 1 (SNED1) | 158.21 | 38 | 3 |
| Q6P3W7 | SCY1-like protein 2 (SCYL2) | 104.33 | 37 | 1 |
| Q06830 | Peroxiredoxin-1 (PRDX1) | 22.32 | 35 | 2 |
| Q5VX52 | Spermatogenesis-associated protein 1 (SPATA1) | 50.50 | 27 | 1 |
| Q02880 | DNA topoisomerase 2-beta (TOP2B) | 184.12 | 27 | 1 |
| P20674 | Cytochrome c oxidase subunit 5A, mitochondrial (COX5A) | 16.92 | 27 | 1 |
| O43150 | Arf-GAP with SH3 domain, ANK repeat and PH domain-containing protein 2 (ASAP2) | 112.84 | 26 | 1 |
| Q9Y2J4 | Angiomotin-like protein 2 (AMOTL2) | 85.94 | 25 | 2 |
| Q99943 | 1-acyl-sn-glycerol-3-phosphate acyltransferase alpha (AGPAT1) | 32.04 | 25 | 1 |

**Table S3.** The list of primers.

| shRNA recovered primers | |
|---|---|
| **Primer** | **Sequence** |
| shRNA-F | 5’-GAGGGCCTATTTCCCATGAT-3’ |
| shRNA-R | 5’-GACGTGAAGAATGTGCGAGA-3’ |
| **qPCR primers** | |
| **Primer** | **Sequence** |
| human NEDD4L-F | 5’-GACATGGAGCATGGATGGGAA-3’ |
| human NEDD4L-R | 5’-GTTCGGCCTAAATTGTCCACT-3’ |
| human AKT1-F | 5’-TCTATGGCGCTGAGATTGTG-3’ |
| human AKT1-R | 5’-TCTTAATGTGCCCGTCCTTG-3’ |
| human AKT2-F | 5’-CGGTTTTATGGTGCAGAGATTG-3’ |
| human AKT2-R | 5’-AGTCAGTGATCTTGATGTGGC-3’ |
| human AKT3-F | 5’-TGTGGATTTACCTTATCCCCTCA-3’ |
| human AKT3-R | 5’-GTTTGGCTTTGGTCGTTCTGT-3’ |
| **TaqMan Gene Expression Assay** | |
| **Gene name** | **Assay ID** |
| CTNNA1 | Hs00944794_m1 |
| CTNNB1 | Hs00355049_m1 |
| JUP | Hs00158408_m1 |
| CDH1 | Hs01023894_m1 |
| VIM | Hs00185584_m1 |
| SOX9 | Hs01001343_g1 |
| POU5F1 | Hs04260367_gH |
| TAZ | Hs00794094_m1 |
| SLUG | Hs00950344_m1 |
| NANOG | Hs04260366_g1 |
| ACTB | Hs01060665_g1 |

**Table S4.** The list of antibodies.

| Antibody | RRID | Imm. Animal | Clone | Cat.# | Vendor | Applications |
|---|---|---|---|---|---|---|
| Anti-NEDD4L | AB_1904063 | Rabbit | -- | 4013 | CST, USA | WB (1:1000) |
| Anti-β-actin | AB_476743 | Mouse | AC-74 | A5316 | Sigma, USA | WB (1:1000) |
| Anti-Flag | AB_259529 | Mouse | M2 | F3165 | Sigma, USA | WB (1:1000) |
| Anti-HA | AB_2770404 | Mouse | 2S8Z1 | AE008 | Abclonal, China | WB (1:1000) |
| Anti-α-Catenin | AB_397592 | Mouse | 5 | 610193 | BD, USA | WB (1:1000) |
| Anti-β-Catenin | AB_397554 | Mouse | 14 | 610153 | BD, USA | WB (1:1000) |
| Anti-γ-Catenin | AB_397648 | Mouse | 15 | 610253 | BD, USA | WB (1:1000) |
| Anti-E-Cadherin | AB_397580 | Mouse | 36 | 610181 | BD, USA | WB (1:1000) |
| Anti-Vimentin | AB_393716 | Mouse | RV202 | 550513 | BD, USA | WB (1:1000) |
| Anti-p-mTOR | AB_10691552 | Rabbit | D9C2 | 5536 | CST, USA | WB (1:1000) |
| Anti-mTOR | AB_330978 | Rabbit | -- | 2972 | CST, USA | WB (1:1000) |
| Anti-tdTomoto | AB_2687917 | Goat | -- | orb182397 | Biorbyt, UK | IF (1:300) |
| Anti-Ki67 | AB_3072239 | Rabbit | SR00-02 | HA 721115 | HUA BIO, China | IF (1:300) |
| Anti-p-4EBP1 | AB_330947 | Rabbit | -- | 9451 | CST, USA | WB (1:1000) |
| Anti-4EBP1 | AB_2097841 | Rabbit | 53H11 | 9644 | CST, USA | WB (1:1000) |
| Anti-p-AKT | AB_2315049 | Rabbit | D9E | 4060 | CST, USA | WB (1:3000) |
| Anti-AKT | AB_915783 | Rabbit | C67E7 | 4691 | CST, USA | WB (1:1000) |
| Anti-AKT | AB_1147620 | Mouse | 40D4 | 2920 | CST, USA | WB (1:1000); IP (1:300) |

| | | | | | | |
|---|---|---|---|---|---|---|
| Anti-Myc | AB_331783 | Mouse | 9B11 | 2276 | CST, USA | WB (1:1000) |
| Anti-PRMT5 | AB_2762092 | Rabbit | -- | A1520 | Abclonal, China | WB (1:3000); IP (1:300) |
| Anti-Ubiquitin | AB_628423 | Mouse | P4D1 | sc-8017 | Santa Cruz, USA | WB (1:200) |
| Anti-WDR77 | AB_2772891 | Mouse | AMC 0495 | A9921 | Abclonal, China | WB (1:1000) |
| Anti-SDMA&MMA | AB_3095615 | Rabbit | -- | PTM-617 | PTM BIO, China | WB (1:500) |
| Anti-AKT1 | AB_3069857 | Rabbit | ST05-09 | ET1609-47 | HUA BIO, China | WB (1:3000); IP (1:300) |
| Anti-AKT2 | AB_3071190 | Rabbit | -- | HA 500091 | HUA BIO, China | WB (1:1000) |
| Anti-Mouse IgG-HRP | AB_631736 | Goat | -- | sc-2005 | Santa Cruz, USA | WB (1:20000) |
| Anti-Rabbit IgG-HRP | AB_631746 | Goat | -- | sc-2004 | Santa Cruz, USA | WB (1:20000) |
| Anti-Rabbit lgG HRP | -- | Mouse | -- | M21006 | Abmart, China | WB (1:1000) |
| Alexa Fluor™ 488-conjugated Donkey anti-rabbit IgG (H+L) | AB_2576217 | Donkey | -- | A21206 | Thermo Fisher Scientific, USA | IF (1:500) |
| Alexa Fluor™ 568-conjugated Streptavidin | AB_2315774 | -- | -- | S11226 | Thermo Fisher Scientific, USA | IF (1:500) |
| Biotinylated horse anti-goat IgG | AB_2336123 | Horse | -- | BA-9500 | Vector Labs, USA | IF (1:300) |

IP, immunoprecipitation assay; IF, immunofluorescence experiments; WB, western blotting.

**Table S5.** The list of sgRNAs.

| sgRNA | Sequence |
|---|---|
| human PRMT5 #1 | 5’-ATGAACTCCCTCTTGAAACG-3’ |
| human PRMT5 #2 | 5’-CCCTTCTCCGTCCCCGAGTT-3’ |
| human AKT1 #1 | 5’-GGGAGTACATCAAGACCTGG-3’ |
| human AKT1 #2 | 5’-ACCGCGTCCTGCAGAACTCC-3’ |
| human AKT2 | 5’-CTCTTCAGCAGGAAGTACCG-3’ |
